# Pathogenesis and natural history of the Bundibugyo species of *Orthoebolavirus* in nonhuman primates

**DOI:** 10.64898/2026.08.10.743937

**Authors:** Karla A. Fenton, Declan D. Pigeaud, Jacquelyn Turcinovic, Abhishek N. Prasad, Krystle N. Agans, Natalie S. Dobias, Rachel O’Toole, Antoinette Lona, Courtney Woolsey, Viktoriya Borisevich, Daniel J. Deer, Joan B. Geisbert, Christopher F. Basler, Robert W. Cross, Thomas W. Geisbert

**Affiliations:** Galveston National Laboratory, University of Texas Medical Branch, Galveston, TX, USA; Department of Microbiology and Immunology, University of Texas Medical Branch, Galveston, TX, USA; Department of Microbiology, Icahn School of Medicine at Mount Sinai, New York, NY, USA

**Author notes:** These authors contributed equally (Karla A. Fenton, Declan D. Pigeaud, Jacquelyn Turcinovic).

## Abstract

The current outbreak of Bundibugyo virus (BDBV) in Africa is a global public health concern particularly as there are no licensed medical countermeasures (MCM). Well characterized animal models that accurately replicate human BDBV infection are needed to develop effective MCM. We exposed 21 cynomolgus monkeys (CM) to BDBV to examine the progression and natural history of BDBV disease (BVD). BVD was more protracted than reported for Ebola and Sudan infection in CM with a lower lethality rate of 67% consistent with lower human BVD mortality rates. IHC and spatial proteomics identified CD209+, CD68+, and/or HLA-DR+ macrophages and dendritic cells as early targets of BDBV. These infected cells frequently colocalized with fibrin and infiltrating MPO+ neutrophils and S100A9+ myeloid-derived suppressor cells, consistent with the development of an active inflammatory response and early coagulopathy. Transcriptomic and proteomic analyses of the circulating immune response correspondingly reflected a cytokine-driven hyperinflammatory state in CM that succumbed to disease. Surviving animals resolved systemic inflammation by the study endpoint; however, BDBV antigen was identified in immune privileged tissues with lesion-associated inflammation aligning with known post-Ebola sequela in humans. This data should assist in identifying weaknesses in the disease course that can be exploited to develop new MCM.

## Introduction

The family *Filoviridae* contains six genera of single-stranded negative-sense RNA viruses two of which, *Orthoebolavirus* and *Orthomarburgvirus,* are significant pathogens of global concern^1,2^. The genus *Orthoebolavirus* consists of six species: *Orthoebolavirus bombaliense*, *Orthoebolavirus bundibugyoense* (Bundibugyo virus; BDBV), Orthoebolavirus restonense*, Orthoebolavirus sudanense* (Sudan virus; SUDV), *Orthoebolavirus taiense*, and *Orthoebolavirus zairense* (Ebola virus; EBOV). BDBV, SUDV, and EBOV have caused lethal infection in humans and nonhuman primates (NHP)^1,2^. Clinical features of orthoebolavirus infection in humans and NHP are the same and include high viremia, an overproduction of cytokines and chemokines, and consumptive coagulopathy, which can lead to multi-organ failure and hypovolemic or septic-like shock.

Before 2013, documented outbreaks of human disease caused by filoviruses occurred sporadically, often separated by years, and geographically contained to the areas of infection in Africa. However, more recently orthoebolaviruses have posed a more significant threat to public health, with several major outbreaks. The 2013-16 West African epidemic of EBOV resulted in 28,600 cases with 11,325 deaths^3^, while the 2018-2020 EBOV outbreak in the Democratic Republic of Congo (DRC) caused 3,481 cases and 2,299 deaths^4^ and is the third largest filovirus outbreak. The current outbreak of BDBV in the Democratic Republic of Congo and Uganda as of June 2026 is projected to exceed 10,000 cases within three months and is already the second largest filovirus outbreak^5^. Historically, most orthoebolavirus outbreaks have been caused by EBOV and SUDV with only two outbreaks prior to 2026 being caused by BDBV^6,7^.

Given that EBOV has caused the most and largest filovirus outbreaks more effort was placed on developing vaccines and treatments for Ebola virus disease (EVD). This emphasis resulted in the development of two licensed vaccines and two licensed human monoclonal antibodies^8^ to combat EVD. While there are no licensed MCM for SUDV a recombinant vesicular stomatitis virus (rVSV)-based vaccine expressing the SUDV glycoprotein, termed VSV-SUDV, was used in a clinical trial during the 2022 SUDV outbreak in Uganda^9^. Remdesivir and the pan-orthoebolavirus human monoclonal antibody (mAb) cocktail MBP134 were also deployed for compassionate use during this 2022 SUDV outbreak^10^. Unfortunately, the development of MCM for Bundibugyo virus disease (BVD) has lagged behind the other orthoebolaviruses.

In response to the current BDBV outbreak, the World Health Organization (WHO) has recommended prioritizing three candidate therapeutics for evaluation in clinical trials among confirmed BVD cases^11^. These include the mAb cocktail MBP134 and the single mAb maftivimab that target the BDBV surface glycoprotein as well as the antiviral remdesivir that targets the BDBV L polymerase. Combination therapy using a monoclonal antibody and remdesivir was also recommended for evaluation. The oral antiviral obeldesivir was recommended for postexposure prophylaxis for contacts of confirmed and probable cases of BVD. Preclinical studies in ferrets and cynomolgus monkeys (CM) showed strong protection against lethal EBOV, SUDV, and BDBV infection when treatment with MBP134 was administered beginning at an advanced stage of disease^12^. However, none of the other recommended therapies have been tested in animals. Remdesivir, which has been used to treat human EBOV infections and was successfully used as a therapy in EBOV-, and SUDV-infected macaques as well as NHPs infected with the closely related Marburg virus (MARV)^13–16^. Interestingly, remdesivir has stronger antiviral activity *in vitro* against BDBV than EBOV^17^. Maftivimab has been shown to have stronger antiviral activity *in vitro* activity against EBOV and SUDV than BDBV^18^ but did not protect NHP against lethal SUDV infection^19^. Combination therapy with a mAb cocktail comparable to MBP134 and remdesivir extended the therapeutic window and provided strong protection to rhesus monkeys infected with MARV and SUDV^14,15^. Obeldesivir provided 80-100% protection in macaques infected with EBOV, SUDV, and MARV when treatment was initiated beginning one day after high dose virus exposure^20–22^.

Regarding vaccines WHO has prioritized the single-dose rVSV-BDBV vaccine that has shown strong protection data against BDBV in nonhuman primates (NHP) both as a preventive vaccine and postexposure treatment^23,24^. The rVSV-BDBV vaccine is based on the same technology as the ERVEBO vaccine that is licensed for EBOV disease. Preclinical studies in cynomolgus monkeys (CM) with a rVSV-EBOV vaccine provided 75% protection against lethal heterologous exposure to BDVD^25^. However, the results of this study are inconclusive as BDBV does not cause uniform lethality in positive control macaques and as clinical disease and viremia were observed in the rVSV-EBOV-vaccinated macaques that survived. In regard to BVD, the WHO has advised that ERVEBO should not be used outside carefully designed research settings^26^.

Animal models have historically played critical roles in the development of effective MCM for human use. While lethal mouse, hamster, and guinea pig models have been developed for EBOV and SUDV by serial adaptation^27^, there are no immunocompetent rodent models for BDBV. We and others have developed ferret models for EBOV, SUDV, and BDBV that employ non adapted wild type viruses^28,29^ and these models should be useful for studying orthoebolavirus infection and assessing the efficacy of MCM. However, the gold standard animal models for orthoebolavirus infection that most accurately reflect human disease are CM and rhesus monkeys^30,31^. We previously showed that BDBV infection of rhesus monkeys resulted in 40% lethality and that survival was associated with early activation of adaptive immunity^32^. In several small studies to assess MCM we and others showed that BDBV infection of CM resulted in ∼ 67-75% lethality^23,30,33^, making CM a more attractive NHP species for assessment of MCM, as smaller study sizes can be employed to achieve statistically significant survival data than with rhesus monkeys. While macaque models have been well characterized for EBOV, SUDV, and MARV there have not been any comparable studies with BDBV. Here, we provide a detailed temporal pathology and natural history study of BDBV-infected CM. Clinical pathology, pathology, virology, transcriptomic, and proteomic findings are reported. Our results improve our understanding of the pathophysiology of BVD and highlight the utility of NHP for testing MCM against BDBV.

## Results

The end stage clinical features of BVD reported during the 2007 and 2012 outbreaks were consistent with features of Sudan virus disease (SVD) and EVD. However, the 2012 BVD outbreak reported a longer incubation period and a longer duration of disease than has been noted for SVD and EVD^7^. This extended disease course appears to recapitulate well with observations in macaques where longer disease courses have been noted for BDV compared to SVD and EVD under near identical test conditions^30^. For example, CM exposed to 1000 PFU of EBOV by the i.m. route succumb between 5 and 8 days after exposure^30,34^, where early studies with BDBV under similar test conditions indicated a window between 9 and 15 days^23–25,33^. Here, our work focused on 1) understanding the progression of early events that lead to lethal disease and 2) shedding additional light on the natural history of BVD in CM using a larger group of animals than previous studies. The prior studies mostly looked at small numbers of animals that served as virus positive controls on MCM studies.

In order to define the progression of events that occur in blood and tissues during BVD infection nine CM were infected with 1000 PFU of BDBV by intramuscular (i.m.) injection. Three animals were euthanized per day at pre-determined timepoints on 2, 4, and 8 days post infection (DPI) (diagrammed in **Figure 1a**). All nine macaques were completely free of overt clinical signs disease up to the 4 DPI timepoint (**Supplementary Table 1**); however, beginning 5 DPI and progressing to the 8 DPI study endpoint, the remaining three macaques began exhibiting signs such as decreased appetite/anorexia (3/3, 100%), depression (2/3, 67%), hunched posture (1/3, 33%), and fever (1/3, 33%).

**Figure 1:**
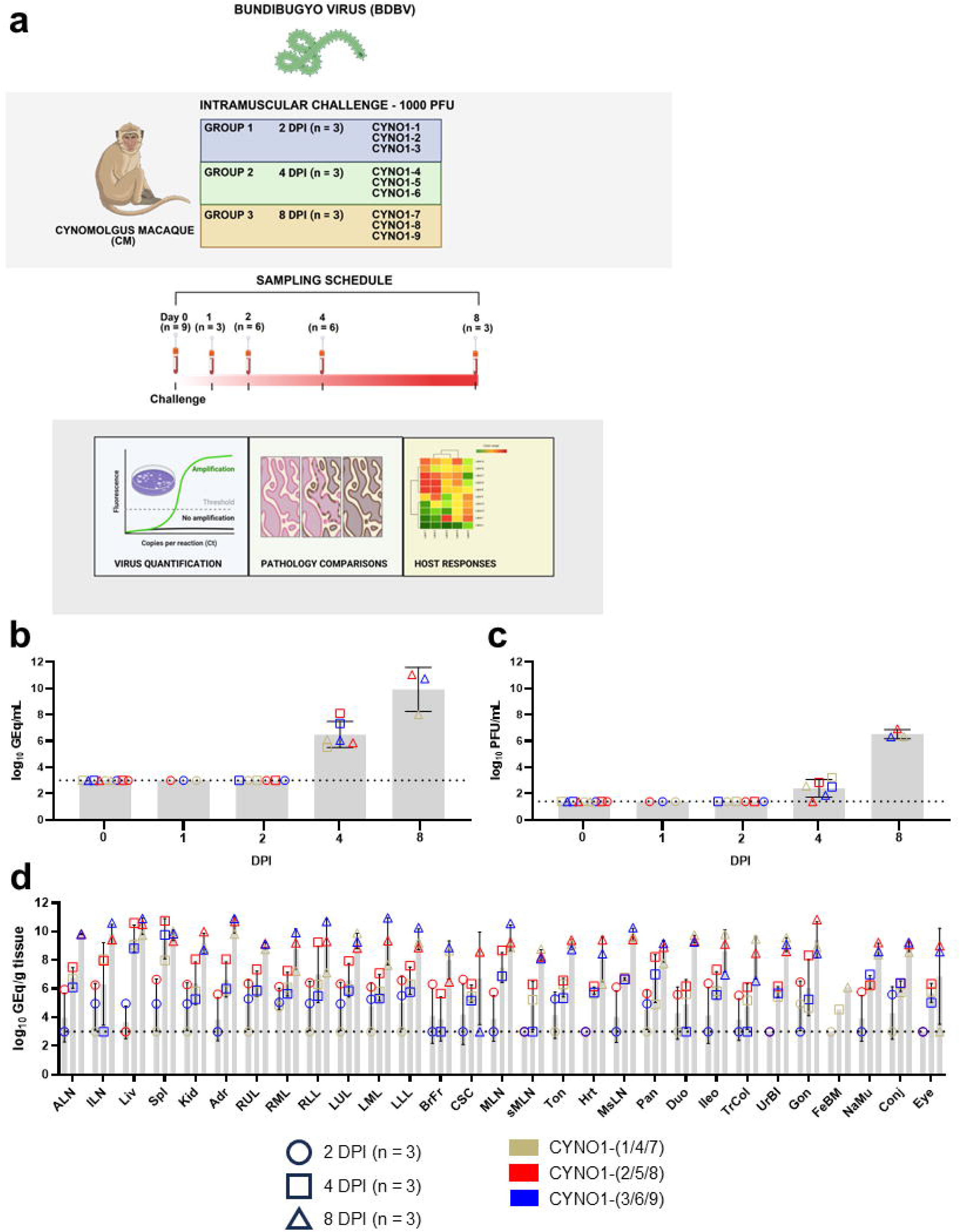
Assessment of viral load in whole blood, plasma, and tissues in cynomolgus macaques from the temporal euthanasia study. **(a)** Diagram of the temporal euthanasia study. This illustration was created in BioRender and is used here by license. **(b)** RT-qPCR detection of BDBV genomic RNA (vRNA) in whole blood from cynomolgus macaques; **(c)** Plaque titration of infectious BDBV in plasma from cynomolgus macaques; **(d)** RT-qPCR detection of BDBV vRNA in tissues from cynomolgus macaques harvested at necropsy. For **(b-d)**, plotted data points represent the mean of two replicate assays, and the bars represent the geometric mean ± geometric SD. Horizontal dashed lines represent the lower limit of quantitation (LLOQ) for the given assay (1000 GEq/mL or 1000 GEq/g tissue for RT-qPCR, 25 PFU/mL for plaque titration). Values below the LLOQ for the RT-qPCR and plaque titration assays are plotted as 999 GEq/mL or gram tissue or 24.9 PFU/mL, respectively. Missing values indicate the tissue was not assayed. Abbreviations: GEq = genome equivalents; ALN = axillary lymph node; ILN = inguinal lymph node; Liv = liver; Spl = spleen; Kid = kidney; Adr = adrenal gland; RUL = right upper lung; RML = right middle lung; RLL = right lower lung; LUL = left upper lung; LML = left middle lung; LLL = left lower lung; BrFr = brain frontal cortex; CSC = cervical spinal cord; MLN = mandibular lymph node; sMnSG = submandibular salivary gland; Ton = tonsil; Hrt = heart; MsLN = mesenteric lymph node; Duo = duodenum; Pan = pancreas; Ile = ileocecal junction; TrCo = transverse colon; UrBl = urinary bladder; Gon = gonads; FeBM = femoral bone marrow; NaMu = nasal mucosa; Conj = conjunctiva; Eye = eye (tissue).

In order to characterize the natural history of BVD a total of 12 CM were infected with 1000 PFU of BDBV by i.m. injection and followed until the predetermined 28 DPI study endpoint or until an animal succumbed to BVD as determined by humane endpoint criteria. Eight of twelve (67%) animals succumbed to BVD 9-17 DPI (mean time to death [MTD] = 13.3 ± 3.4 DPI), while four animals survived to the 28 DPI predetermined study endpoint (**Figure 2a**). All 12 animals developed overt clinical signs of disease (**Supplementary Table 2**), including some or all of the following: fever (6/12, 50%), decreased appetite and/or anorexia (12/12, 100%), depression/reduced activity (12/12, 100%); petechial rash (8/12, 67%), and hunched posture (11/12, 92%). Morbidities such as ataxia (2/12, 17%) and dyspnea (3/12, 25%) were less commonly observed and correlated with a fatal clinical outcome.

**Figure 2:**
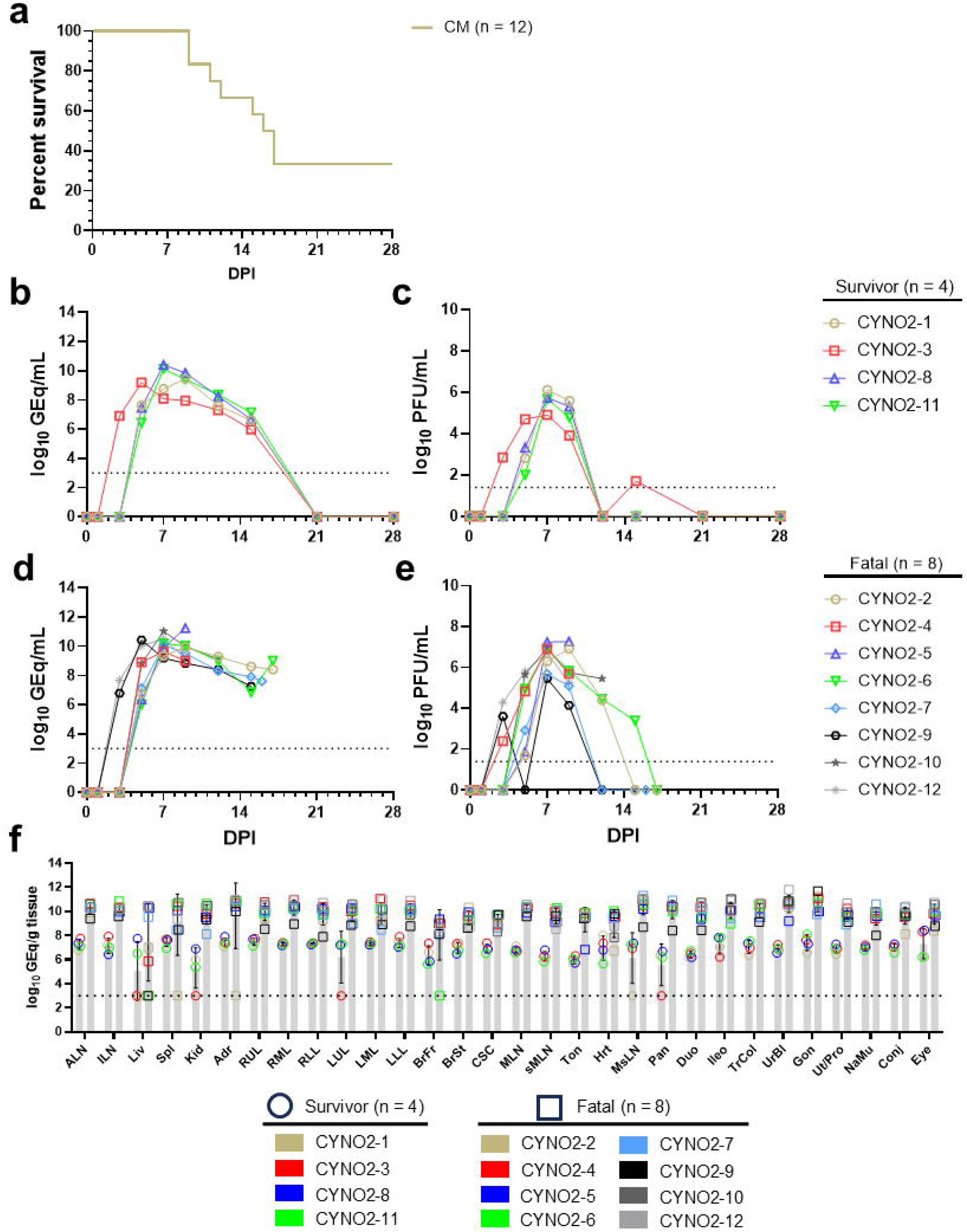
Survival analysis and assessment of viral load in whole blood, plasma, and tissues in cynomolgus macaques from the natural history study. **(a)** Kaplan-Meier survival curve of cynomolgus macaques challenged with BDBV; **(b)** RT-qPCR detection of BDBV genomic RNA (vRNA) in whole blood from cynomolgus macaques that survived challenge with BDBV; **(c)** Plaque titration of infectious BDBV in plasma from cynomolgus macaques that survived challenge with BDBV **(d)** RT-qPCR detection of BDBV vRNA in whole blood from cynomolgus macaques that developed lethal disease following challenge with BDBV; **(e)** Plaque titration of infectious BDBV in plasma from cynomolgus macaques that developed lethal disease following challenge with BDBV; **(f)** RT-qPCR detection of BDBV vRNA in tissues from cynomolgus macaques harvested at necropsy. For **(b-f)**, plotted data points represent the mean of two replicate assays. Horizontal dashed lines represent the lower limit of quantitation (LLOQ) for the given assay (1000 GEq/mL or 1000 GEq/g tissue for RT-qPCR, 25 PFU/mL for plaque titration). For **(f)**, bars represent the geometric mean ± geometric SD. Values below the LLOQ for the RT-qPCR assay are plotted as 999 GEq/gram tissue. Missing values indicate the tissue was not assayed. Abbreviations: GEq = genome equivalents; ALN = axillary lymph node; ILN = inguinal lymph node; Liv = liver; Spl = spleen; Kid = kidney; Adr = adrenal gland; RUL = right upper lung; RML = right middle lung; RLL = right lower lung; LUL = left upper lung; LML = left middle lung; LLL = left lower lung; BrFr = brain frontal cortex; BrSt = brain stem; CSC = cervical spinal cord; MLN = mandibular lymph node; sMnSG = submandibular salivary gland; Ton = tonsil; Hrt = heart; MsLN = mesenteric lymph node; Duo = duodenum; Pan = pancreas; Ile = ileocecal junction; TrCo = transverse colon; UrBl = urinary bladder; Gon = gonads; Ut/Pro = uterus/prostate; NaMu = nasal mucosa; Conj = conjunctiva; Eye = eye (tissue).

### Clinical pathology

In the temporal euthanasia study, mild perturbations to baseline hematological and serum biochemical markers were observed up to the 4 DPI timepoint, which included monocytopenia (4/9, 44%), and granulocytosis (6/9, 67%). Only a single animal (CYNO1-1) exhibited a significant increase in a marker of hepatic insult at 1 and 2 DPI (AST, < 5-fold increase from baseline), and a different animal (CYNO1-5) exhibited lymphocytopenia and elevated CRP at 4 DPI (**Supplementary Table 1**). By 8 DPI, the remaining three macaques exhibited marked deviations in hematological profiles, including lymphocytopenia (3/3, 100%), thrombocytopenia (3/3, 100%), and granulocytosis (2/3, 67%), and more pronounced increases in serum markers of hepatic and renal injury such as ALT (3/3, 100%), AST (3/3, 100%), ALP (3/3, 100%), and CRP (2/3, 67%). Animals in the natural history study followed a similar temporal progression in the severity of disruption to baseline hematological values, most of which gradually returned to baseline values in surviving animals by 15 DPI (**Supplementary Table 2, Supplementary Figure 1**). A panel of analytes that showed the greatest perturbations from baseline and that are known markers of filovirus disease were selected for statistical analysis. There was no significant difference in fold-change values from baseline in macaques that survived challenge versus those that succumbed to lethal disease (**Supplementary Figure 1**).

### Virology

We assessed levels of circulating and BDBV RNA and infectious BDBV by RT-qPCR and plaque titration, respectively. For the temporal euthanasia study, we did not detect circulating BDBV RNA or infectious BDBV in any animal at 2 DPI (**Figure 1b, c**). By 4 DPI all remaining macaques had moderate levels of circulating BDBV RNA (5.51-8.10 log_10_ GEq/ml) while 5/6 animals had low levels of circulating infectious BDBV (1.88-3.22 log_10_ PFU/ml) that progressed by 8 DPI, where all remaining macaques had moderate to high levels of circulating BDBV RNA (8.00-11.03 log_10_ GEq/ml) and infectious BDBV (6.31-6.92 log_10_ PFU/ml) (**Figure 1b, c**). For the natural history study, we did not detect circulating BDBV RNA or infectious BDBV in any animal until 3 DPI when 3/12 macaques had moderate levels of circulating BDBV RNA (6.79-7.68 log_10_ GEq/ml) and the same three animals and one additional macaque had detectable infectious BDVD at this time point (2.40-4.30 log_10_ PFU/ml) (**Figure 2b-e**). By 5 DPI, all animals had detectable levels of circulating BDBV RNA (6.01-7.42 log_10_ GEq/ml) and 11/12 had detectable infectious virus (1.70-5.80 log_10_ PFU/ml). BDBV RNA and infectious virus were detected in all animals by 7 DPI with peak levels occurring in most animals at 7 or 9 DPI (8.09-11.03 log_10_ GEq/ml at 7 DPI and 7.95-11.25 log_10_ GEq/ml at 9 DPI; 4.90-7.25 log_10_ PFU/ml at 7 DPI and 3.90-7.27 log_10_ PFU/ml at 9 DPI). Circulating BDBV RNA and infectious virus was cleared in all four surviving macaques by 21 DPI. In order to determine the effect of circulating viral load on survival outcome, we compared the peak vRNA abundance and circulating viremia between macaques that survived challenge versus those that developed lethal disease. For statistical analysis, we utilized data from 9 historical positive control cynomolgus macaques from previous studies that were challenged with the exact same virus stock, challenge dose, and inoculation route (8/9 [89%] fatal, MTD = 12.0 DPI). There was no significant difference in the peak abundance of circulating vRNA between surviving macaques and those that developed lethal disease; however, peak circulating infectious virus titers were significantly lower in macaques that survived challenge versus those that did not (**Supplementary Figure 2a,b**, p = 0.003, Mann-Whitney U-test,).

Tissues were collected at necropsy from all animals and assessed for the presence of BDBV RNA by RT-qPCR. For the temporal euthanasia study, the number of tissues with detectable BDBV RNA and the amount of BDBV RNA detected increased with each time point with nearly all tissues from all animals having moderate to high levels of BDBV RNA by 4 DPI with liver and spleen generally having the highest viral loads (7.98-10.75 log_10_ GEq/g tissue) at that time point (**Figure 1d**). In the natural history study, high levels of BDBV RNA (> 9.00 log_10_ GEq/g tissue) were detected in most tissues of all eight animals that succumbed to BVD (**Figure 2f**). Low to moderate levels of BDBV (< 8.5 log_10_ GEq/g tissue) were detected in some tissues of all of the four surviving macaques. In contrast, previous studies by our group have never detected infectious orthebolaviruses or MARV at these low to moderate levels^20–22^. Evidence of residual vRNA in the absence of infectious virus is not unexpected in recovered animals and is likely due to the presence of ongoing immune clearance mechanisms (that is, neutralizing antibodies, cellular immunity and so on) and has been documented by us in arenavirus vaccine and treatment studies^35,36^ and by others for a number of other virus infection models^37^.

### Assessment of BDBV GP-specific IgG binding titers

Enzyme-linked immunosorbent assays (ELISA) were performed on sera collected from all macaques from the natural history study at pre-determined timepoints (0, 7, 9, 15, and 28 DPI), or at the terminal timepoint for the animal, and binding antibody titers were determined using the endpoint dilution method. BDBV GP-specific IgG was first detected at 12 DPI in a single terminal subject that was assayed; all other surviving macaques had reciprocal dilution titers ranging from 1600 to 6400 by 15 DPI (**Supplementary Figure 3**). Titers in animals that survived to the 28 DPI study endpoint ranged from 6400 to 25600. Statistical comparison was performed on IgG titers from animals that survived challenge versus those that succumbed only for the 15 DPI timepoint; the difference was not significant (p = 0.43, Mann-Whitney U-test).

### Gross lesions and histopathology

For the temporal euthanasia study, the only pronounced gross findings were mild multicentric lymphadenomegaly starting at 4 DPI and continuing into 8 DPI and mild hepatic reticulation at 8 DPI. Histologic evaluation of tissues from animals euthanized at 2 DPI (CYNO1-1, CYNO1-2, and CYNO1-3) revealed no appreciable microscopic lesions and no detectable BDBV antigen by immunohistochemistry (IHC) in any examined tissues (**Figure 3a,d,g**). By 4 DPI, minimal lesions and limited BDBV antigen distribution were evident in all three animals (CYNO1-4, CYNO1-5, and CYNO1-6). Lesions were restricted primarily to vascular compartments, lymphoid tissues, and liver (**Supplementary Table 3**). Mild expansion of peripheral lymphoid tissues by leukocytes (lymphoid histiocytosis), increased leukocytes within hepatic sinusoids (sinusoidal leukocytosis) were observed and corroborated the finding of multicentric lymphadenomegaly from gross examination. Viral antigen was localized predominantly to scattered mononuclear cells morphologically consistent with histiocytes, macrophages, and/or dendritic cells within hepatic sinusoids and lymphoid tissues, including the gut associated lymphoid tissues (GALT) (**Figure 3b,e,h**), indicating an early tropism for cells of the mononuclear phagocyte system.

**Figure 3:**
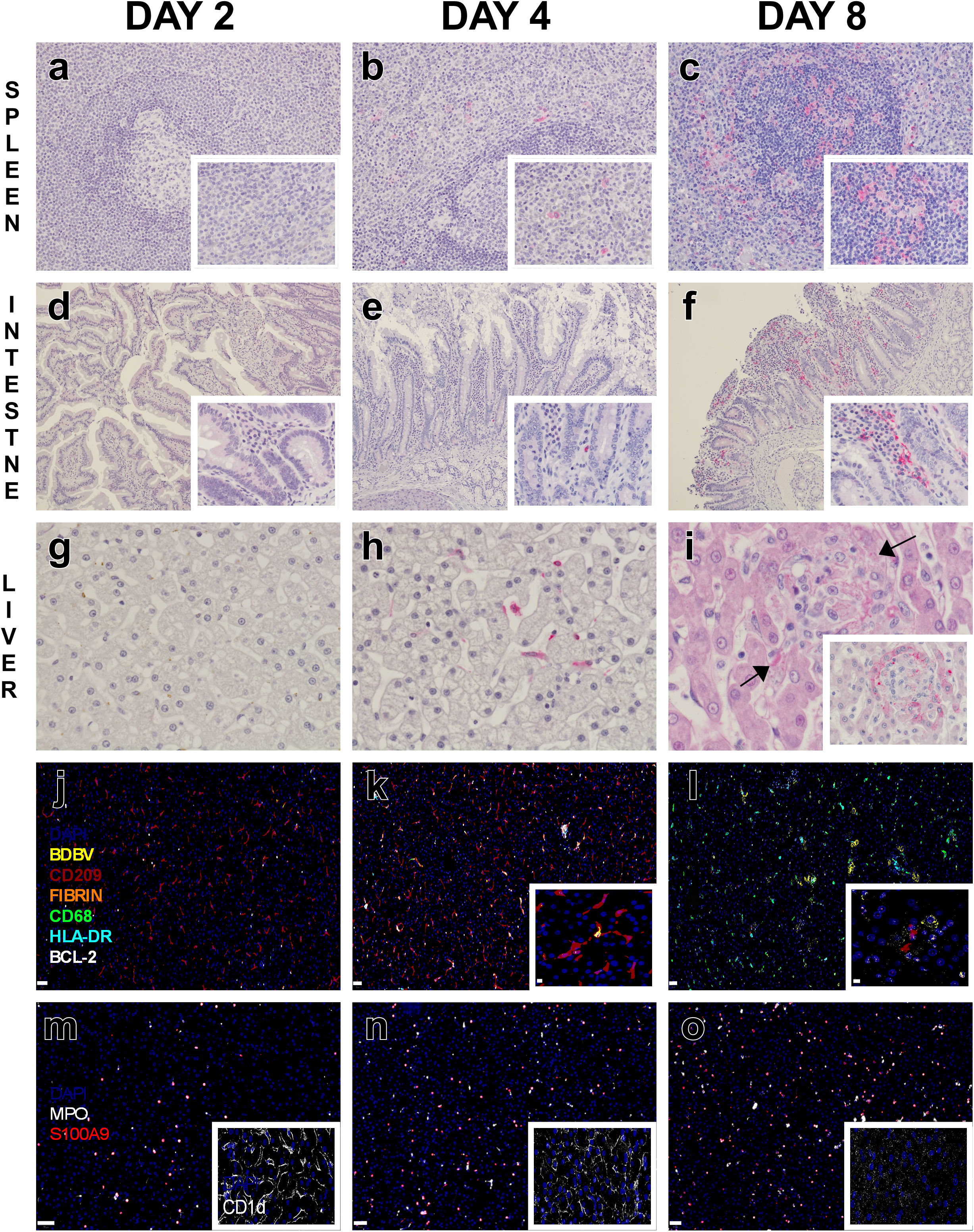
Temporal histopathology, immunohistochemistry (IHC), and multiplex immunofluorescence (mIF) of tissues from BDBV-infected NHPs. Representative hematoxylin and eosin (H&E), IHC images for anti-BDBV GP antibody (red), and mIF in cynomolgus macaques (CM) challenged with BDBV and euthanized on 2 DPI (**a,d, g, j, m**), 4 DPI (**b,e,h,k,n**), or 8 DPI (**c,f,i,l,o**). Images captured at 10x magnification (**d-f**), 20x magnification (**a-c**), 40x magnification (**g,h)**, and 60x magnification (**a-c** insets, **d-f** inset, i**, i** inset,). Images representing spleen (**a-c**), intestine (**d-f**), liver (**g-o**). **a)** Spleen, 20x, CYNO1-2, Day 2: No appreciable IHC positive (red) labeling. Inset-60x higher magnification. **b)** Spleen, 20x, CYNO1-5, Day 4: Scattered mononuclear cells within the red pulp were IHC positive (red). Inset-60x higher magnification. **c)** Spleen, 20x, CYNO1-9, Day 8: Moderate numbers of mononuclear cells within the red and white pulp were IHC positive (red). Inset-60x higher magnification. **d)** Intestine, 10x, CYNO1-2, Day 2: No appreciable IHC positive (red) labeling. Inset-60x higher magnification. **e)** Intestine, 10x, CYNO1-5, Day 4: Minimal expansion of the lamina propria by lymphohistiocytic inflammatory infiltrates in which scattered mononuclear cells were IHC positive (red). Inset-60x higher magnification. **f)** Intestine, 10x, CYNO1-7, Day 8: Moderate villous blunting, fusion, and epithelial sloughing, with expansion of the lamina propria by lymphohistiocytic inflammatory infiltrates in which numerous mononuclear cells were IHC positive (red). Inset-60x higher magnification. **g)** Liver, 40x, CYNO1-2, Day 2: No appreciable IHC positive (red) labeling. **h)** Liver, 40x, CYNO1-5, Day 4: Scattered mononuclear cells within the hepatic sinusoids were IHC positive (red). **i)** Liver, 40x, CYNO1-7, Day 8: Focal hepatocellular degeneration and necrosis with clustered mononuclear cells expanding the hepatic sinusoids with prominent cytoplasmic amorphous eosinophilic inclusion bodies within hepatocytes consistent with viral inclusions (arrows). Inset-60x, IHC, Colocalization of IHC positivity and inclusion bodies within hepatocytes. **j)** Day 2, Cy CYNO1-2. Abundant CD209+ (Red) cells and few scattered CD209+ CD68+ (Green) cells, few scattered BCL-2+ (White) T-cells, within liver sinusoids. No labeling for BDBV (Yellow), Fibrin (Orange), HLA-DR (Light blue). DAPI (Blue) nuclear stain. Scale bar 50um. **k)** Day 4, Cy CYNO1-5. Moderate numbers of CD209+, CD68+, HLA-DR+, BDBV+, BCL-2+, Fibrin+ cells colocalized within liver sinusoids. DAPI (Blue) nuclear stain. Scale bar 50um. Inset, higher magnification of CD209+, BDBV+ cells. Scale bar 5um. **l)** Day 8, Cy CYNO1-9. Moderate numbers of BDBV+ hepatocytes and CD68+ cells, sparse CD209+, HLA-DR+, BDBV+ cells. DAPI (Blue) nuclear stain. Scale bar 50um. Inset, higher magnification of CD209+, BDBV+ cells. Scale bar 5um. **m)** Day 2, Cy CYNO1-2. Scattered MPO+ (White) and S100A+ (Red) cells within liver sinusoids. DAPI (Blue) nuclear stain. Scale bar 50um. Inset: Day 2, Cy CYNO1-2. Abundant CD1d (White) labeling of hepatocytes. DAPI (Blue) nuclear stain. **n)** Day 4, Cy CYNO1-5. Moderate numbers of MPO+ (White) and S100A+ (Red) cells within liver sinusoids. DAPI (Blue) nuclear stain. Scale bar 50um. Inset: Day 4, Cy CYNO1-5. Moderate CD1d (White) labeling of hepatocytes. DAPI (Blue) nuclear stain. **o)** Day 8, Cy CYNO1-9. Marked numbers of MPO+ (White) and S100A+ (Red) cells within liver sinusoids. DAPI (Blue) nuclear stain. Scale bar 50um. Inset: Day 8, Cy CYNO1-9. Disrupted CD1d (White) labeling of hepatocytes. DAPI (Blue) nuclear stain.

By 8 DPI, disease progression was characterized by marked expansion of lesions within lymphoid organs and liver, accompanied by widespread systemic dissemination of BDBV-positive mononuclear cells. In 2/3 macaques, BDBV antigen was readily identified in highly vascularized organs, including the lung, heart, kidney, urinary bladder, endocrine glands, and exocrine tissues, uterus, prostate, brain, and eye (**Supplementary Figure 4a-o**). Lymphoid lesions were most prominent. Peripheral lymph nodes exhibited marked sinus expansion by histiocytes, with multifocal clusters to sheets of antigen-positive mononuclear cells. In the spleen, red pulp hypercellularity was associated with increased leukocyte infiltration, while white pulp contained areas with tingible-body macrophages and karyorrhectic debris, indicative of lymphocytolysis and apoptosis. Numerous BDBV-positive mononuclear cells were distributed throughout the red pulp and occasionally concentrated adjacent to areas of lymphoid depletion (**Figure 3c**). Mucosal-associated lymphoid tissues were similarly affected. Gastrointestinal involvement was observed throughout all examined segments of the alimentary tract and was characterized by expansion of the lamina propria by mononuclear inflammatory cells, many of which were strongly positive for BDBV antigen. Variable villous blunting and fusion accompanied the inflammatory infiltrates which impair absorption and potentially lead clinically to gastrointestinal distress (**Figure 3f**. Although intestinal epithelial cells were generally IHC negative, frequent epithelial sloughing prevented definitive assessment of epithelial infection and raised the possibility that infected epithelium and inflammatory cells within the mucosa could contribute to luminal viral shedding. Within the tonsil, clusters of BDBV-positive mononuclear cells were distributed throughout lymphoid follicles, along the epithelial interface, and rare BDBV-positive epithelial cells within the overlying mucosa (**Supplementary Figure 4a**).

Hepatic lesions progressed from sinusoidal leukocytosis to multifocal periportal to random hepatitis with aggregates of BDBV-positive mononuclear cells and hepatocytes (**Figure 3i inset**). Multifocal centrilobular granulocytic and histiocytic vasculitis was present in one macaque, with BDBV antigen present in the infiltrating inflammatory cells (CYNO1-9). Prominent cytoplasmic amorphous eosinophilic inclusion bodies consistent with viral inclusions were observed in clusters of hepatocytes and colocalized with BDBV antigen (**Figure 3i**), indicating productive infection of hepatocytes at later stages of disease.

Pulmonary lesions consisted of mild to moderate interstitial pneumonia in all 8 DPI animals, with expansion of alveolar septa by infiltrating leukocytes, a subset of which were BDBV-positive (**Supplementary Figure 4b**). Two macaques (CYNO1-7 and CYNO1-9) had BDBV-positive interstitial cells within the myocardium in the absence of significant inflammation (**Supplementary Figure 4c**). Renal lesions were minimal and consisted of mild interstitial inflammation and degeneration of proximal convoluted tubules. BDBV antigen was present within inflammatory cells of the interstitium and within leukocytes occupying glomerular capillary tufts (**Supplementary Figure 4d**). In the urinary bladder, BDBV-positive inflammatory cells were present within the subepithelial stroma, and rare transitional epithelial cells demonstrated immunoreactivity (**Supplementary Figure 4e**).

Extensive dissemination of BDBV antigen was observed throughout the reproductive tract. In females, BDBV-positive mononuclear cells were identified within the ovarian medulla and among cells associated with the theca interna and externa of follicles at multiple stages of maturation, including atretic follicles (**Supplementary Figure 4f**). The infundibular stroma was expanded by BDBV-positive inflammatory cells, and multifocal vascular endothelial labeling was observed within small- to medium-caliber vessels (**Supplementary Figure 4g**). The uterus contained dense bands of BDBV-positive stromal cells immediately subjacent to the glandular epithelium (**Supplementary Figure 4h**). In males, BDBV-positive mononuclear cells were present within the highly vascularized interstitium of the epididymis and testis and within inflammatory nodules of the prostate (**Supplementary Figure 4i, j**). Rare BDBV-positive cells within the glandular epithelial cells were observed within prostatic acini (**Supplementary Figure 4j**). The presence of BDBV antigen within reproductive tract epithelia and adjacent stromal tissues suggests a potential mechanism for viral shedding in reproductive secretions.

Endocrine tissues demonstrated widespread involvement, with antigen-positive cells identified within pancreatic islets, thyroid parafollicular cells, adrenal cortex, and pituitary gland (**Supplementary Figure 4k**). BDBV-positive cells were present within the conjunctiva, and mucosal-associated lymphoid tissues. In the skin, antigen-positive cells were observed within connective tissue sheaths surrounding hair follicles (**Supplementary Figure 4l**), and inflammatory infiltrates surrounding mammary ducts frequently contained viral antigen (**Supplementary Figure 4m**). In some animals, sebaceous glands of the haired skin also demonstrated positive labeling (**Supplementary Figure 4n**). Multifocal periductal inflammatory infiltrates containing BDBV-positive cells were observed within the submandibular salivary gland. Within the central nervous system, BDBV-positive cells were detected in the choroid plexus of one animal (**Supplementary Figure 4o**). Ocular tissues also contained BDBV antigen, including cells within the ciliary body and choroid.

To further characterize cell types involved in the BVD CM model, multiplex immunofluorescence (mIF) was performed on representative liver sections. mIF revealed progressive changes in inflammatory cell populations and BDBV distribution. At 2 DPI, abundant CD209 (DC-SIGN)+ cells, consistent with dendritic cell lineage, exhibited absent to minimal co-expression of BDBV antigen, CD68 (a macrophage/lysosomal marker), HLA-DR (MHC class II antigen), MPO (myeloperoxidase, a neutrophil/granulocytic enzyme), and S100A9 (a calcium-binding protein associated with inflammatory myeloid cells). (**Figure 3 j,m**). BCL-2 (B-cell lymphoma 2, an anti-apoptotic protein) + T-cell populations were consistently present throughout the study and showed no appreciable temporal changes. By 4 DPI, moderate numbers of BDBV+CD209+ and BDBV+CD209+CD68+HLA-DR+ cells were detected within hepatic sinusoids, indicating infection of hepatic mononuclear phagocytes and antigen-presenting cells. Rare BDBV+CD209+ cells also co-expressed BCL-2. These infected cells frequently colocalized with sparse fibrin deposits and increased MPO+ and S100A9+ infiltrates, consistent with the onset of hepatic inflammation and early coagulopathy (**Figure 3k,n).** At 8 DPI, BDBV antigen was abundant throughout the liver, with extensive hepatocyte labeling and only occasional BDBV-positive CD209+ or HLA-DR+ cells, indicating viral dissemination beyond the initial immune cell targets. Concurrently, CD209+ cells were markedly reduced, whereas CD68+ and CD68+HLA-DR+ macrophage populations increased, accompanied by a substantial influx of MPO+ and S100A9+ myeloid cells, consistent with severe hepatic inflammation (**Figure 3 l,o**). CD1d expression was readily detected on hepatocytes at 2 and 4 DPI but was markedly diminished by 8 DPI, coinciding with extensive hepatocellular degeneration and necrosis (**Figure 3 m-o, insets**).

For the natural history study, gross examination demonstrated widespread multisystemic disease in animals that succumbed to infection. Hepatitis was observed in all eight fatal cases (**Figure 4d**: CYNO2-10). Multicentric lymphadenomegaly and splenomegaly were present in 7/8 animals (CYNO2-2, CYNO2-4, CYNO2-5, CYNO2-7, CYNO2-9, CYNO2-10, CYNO-2-12), highlighting the lymphoid system and liver as major targets of BDBV infection. Additional gross findings included gastroenteritis (CYNO2-2, CYNO2-4, CYNO2-5, CYNO2-10 [**Figure 4i**], and CYNO2-12), petechial, ecchymotic, and/or macular cutaneous rash (CYNO2-2, CYNO2-4, CYNO2-5, CYNO2-6, CYNO2-10 [**Figure 4n**], and CYNO2-12), interstitial pneumonia (CYNO2-6, CYNO2-7, CYNO2-9, and CYNO2-12), meningeal congestion of the brain (CYNO2-2, CYNO2-6, CYNO2-7, CYNO2-9, CYNO2-10, and CYNO2-12), adrenomegaly (CYNO2-2, CYNO2-5, and CYNO2-12), and ascites (CYNO2-9). Histologic evaluation of tissues from the eight CM that succumbed to BVD corroborated gross findings and expanded upon lesions previously identified during the temporal progression of disease. The most severe lesions consisted of widespread fibrin deposition, necrosis and apoptosis affecting large populations of parenchymal and immune cells, predominantly within lymphoid tissues, mucosal-associated lymphoid tissues (MALT), liver, pancreas, and, less frequently, haired skin and mammary gland. BDBV antigen was widely distributed within mononuclear cells, endothelial cells, and parenchymal epithelial cells and was consistently associated with sites of inflammation and tissue injury (**Supplementary Table 4**). Animals that succumbed later in disease progression (CYNO2-2, CYNO2-6, CYNO2-7, and CYNO2-9; 15-17 DPI) exhibited reduced lesion severity and BDBV antigen burden within the lung, peripheral lymphoid tissues, and liver relative to earlier fatalities; however, sustained lesion severity and/or antigen distribution were observed within the brain, pancreas, gastrointestinal tract, urogenital, endocrine organs, heart, haired skin, and eye. In contrast, surviving animals (CYNO2-1, CYNO2-3, CYNO2-8, and CYNO2-11) displayed only minimal residual lesions consisting of BDBV-positive cells in the brainstem, mild lymphocytic infiltrates within the liver and kidney and ischemic dermatitis with necrosis of the tail skin in CYNO2-3. No additional lesions or viral antigen labeling attributable to BDBV were identified in surviving animals.

**Figure 4:**
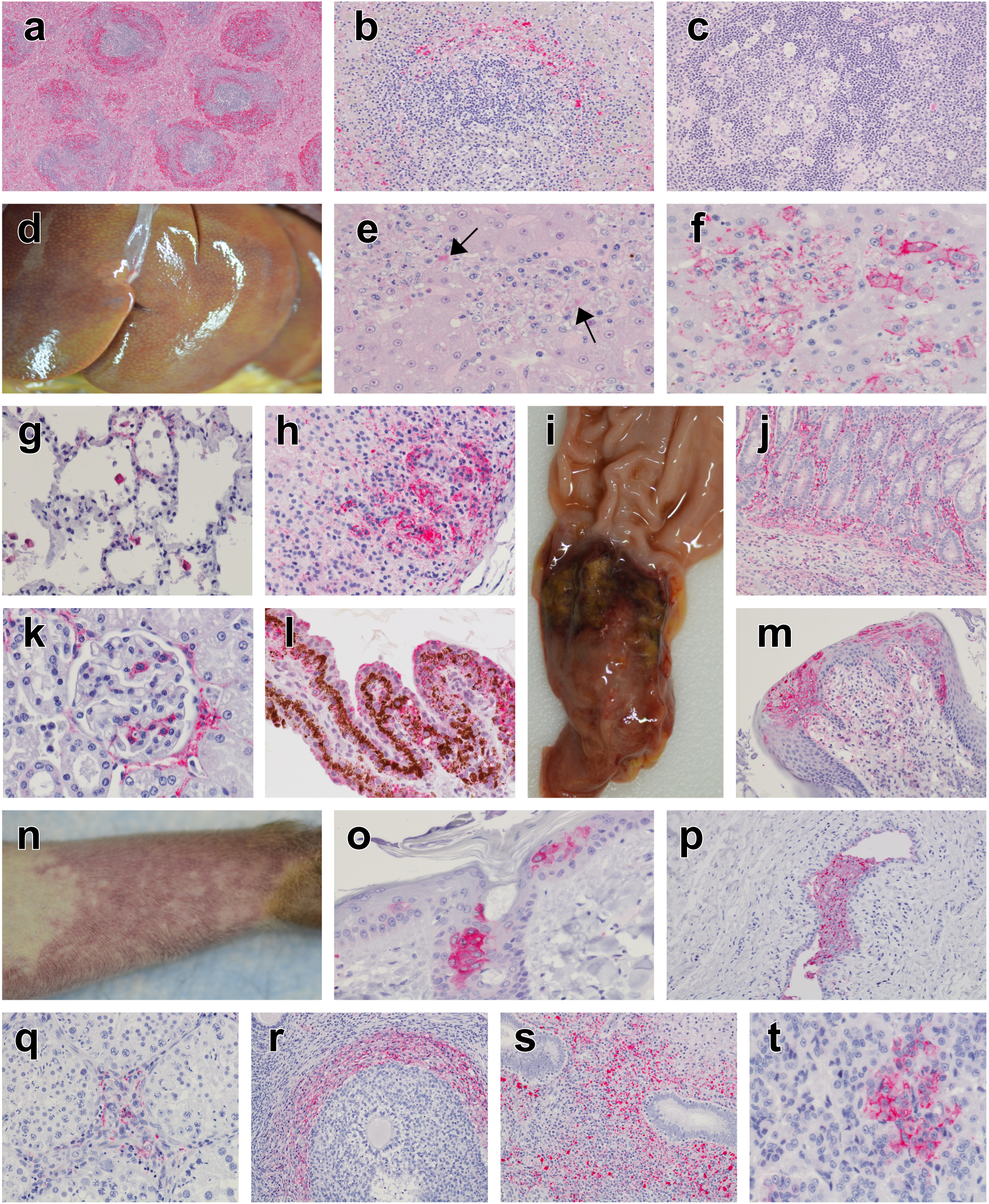
BDBV natural history of gross pathology, histopathology and immunohistochemistry (IHC) of tissues from the lethally-infected NHP. Representative gross pathology, H&E and IHC images for anti-BDBV GP antibody (red) in cynomolgus macaques (CM) infected with BDBV. Images captured at 4x magnification (a), 20x magnification (b,c,j,m,p,r,s), 40x magnification (e,f,g,h,l,o,q), and 60x magnification (k,t). **a)** Spleen, 4x, H&E, CYNO2-5, Day 11: Lymphocytolysis with hemorrhage into white pulp and fibrin accumulation in the red pulp **b)** Spleen, 20x, CYNO2-4, Day 9: Mononuclear cells within the red and white pulp were IHC positive (red). **c)** Lymph Node (Mandibular), 20x, CYNO2-5, Day 11: Tingable body macrophages within germinal center **d)** Liver, gross, CYNO2-10, Day 12: Hepatitis **e)** Liver, 40x, CYNO2-12, Day 9: Necrotizing hepatitis with intracytoplasmic amorphous eosinophilic inclusion bodies within hepatocytes consistent with viral inclusions (arrows). **f)** Liver, 40x, CYNO2-12, Day 9: Hepatocytes and mononuclear inflammatory were IHC positive (red). **g)** Lung, 40x, CYNO2-4, Day 9: Minimal expansion of alveolar septa with mononuclear cells and increased alveolar macrophages of which some are IHC positive (red). **h)** Adrenal Gland, 40x, CYNO2-4, Day 9: IHC positive adrenal cortical cells (red). **i)** Small Intestine (duodenum), gross, 1302755, Day 12: Ulcerative enteritis **j)** Intestine, 20x, CYNO2-12, Day 9: Minimal expansion of the lamina propria by lymphohistiocytic inflammatory infiltrates in which mononuclear cells were IHC positive (red). **k)** Kidney, 60x, CYNO2-4, Day 9: IHC positive mononuclear cells within the capillary bed of the glomerular tuff and surrounding interstitial spaces. **l)** Eye, 40x, CYNO2-9, Day 15: IHC positive epithelium and mononuclear cells of the ciliary body **m)** Tongue, 20x, CYNO2-12, Day 9: IHC positive epithelium and mononuclear cells within the subepithelial stroma **n)** Antebrachium, gross, CYNO2-10, Day 12: Ecchymotic rash **o)** Haired skin of the face, CYNO2-7, Day 16: IHC positive epithelium of the epidermis and hair follicle **p)** Mammary gland, 20x, CYNO2-7, Day 16: IHC positive epithelium and luminal contents of the ducts of the mammary gland **q)** Testis, CYNO2-10, Day 12: IHC positive cells within interstitium of the testis **r)** Ovary, CYNO2-4, Day 9: IHC positive thecal cells **s)** Uterus, CYNO2-9, Day 15: IHC positive uterine stromal cells and inflammatory mononuclear cells **t)** Pituitary gland, CYNO2-4, Day 9: IHC positive cells of the anterior pituitary (adenohypophysis)

Consistent with the temporal pathology findings described, lymphoid tissues, MALT, and liver represented the principal target organs of fatal BDBV infection. Lymphoid tissues exhibited severe lymphocytolysis, hemorrhage, fibrin deposition, and abundant viral antigen, particularly within the spleens of CYNO2-2, CYNO2-5 (**Figure 4a,b**), CYNO2-6, CYNO2-10, and CYNO2-12, with similar but less severe changes in CYNO2-4, CYNO2-7, and CYNO2-9. Lymph nodes, tonsils, and other MALT tissues contained tingible body macrophages and widespread BDBV-positive mononuclear and endothelial cells (**Figure 4c**). Gastroenteritis was observed in all fatal cases with BDBV-positive inflammatory cells and occasional enterocytes (**Figure 4j**), while glossitis was observed in CYNO2-4, CYNO2-5, CYNO2-10, and CYNO2-12 (**Figure 4m**). Hepatic lesions progressed to multifocal hepatitis with abundant BDBV-positive mononuclear cells, hepatocytes, and endothelial cells, frequently accompanied by viral inclusion bodies (**Figure 4e,f**).

Although less extensive than lymphoid and hepatic involvement, CNS lesions were identified in all succumbed animals and were concentrated in or near regions with highly vascular structures lacking a conventional blood-brain barrier. Lymphohistiocytic choroid plexitis was present in all succumbed animals, while meningitis and perivascular cuffing within the neuroparenchyma were additionally observed in CYNO2-2, CYNO2-6, and CYNO2-12. Similar inflammatory lesions and BDBV-positive cells were consistently identified within the anterior pituitary gland, and four animals also demonstrated involvement of the posterior pituitary (**Figure 4t**). Among survivors, only one animal (CYNO2-1) contained rare BDBV-positive cells near the area postrema of the brainstem, a circumventricular organ located within the floor of the fourth ventricle (**Supplementary Figure 5o**). All remaining survivors lacked significant lesions or BDBV antigen (**Supplementary Figure 5a-n,p,q**).

mIF was employed to further characterize the pathology of neuronal tissues of CYNO2-6 at 17 DPI. BDBV antigen and inflammation concentrated within the choroid plexus and perivascular regions of the thalamus within the examined tissue sections. BDBV antigen was detected within IBA-1+ macrophages/microglia, including activated IBA-1+/HLA-DR+ cells, as well as within choroid plexus epithelial cells. These lesions were accompanied by infiltrating MPO+/S100A9+ myeloid cells and minimal fibrin deposition, consistent with an active inflammatory response (**Figure 5a**). Within the thalamus, BDBV-positive macrophages and activated CD68+/HLA-DR+ inflammatory cells were concentrated around blood vessels exhibiting marked loss of Claudin-5 expression and increased Aquaporin-4 labeling in adjacent astrocytic endfeet. These findings provide strong evidence of blood-brain barrier disruption characterized by endothelial tight junction loss, vascular leakage, and edema. Endothelial cells not associated with inflammation also exhibited increased HLA-DR expression, indicating endothelial activation (**Figure 5b**). Areas of perivascular inflammation were further associated with reduced neuronal density and loss of NeuN expression in MAP2+ neurons, consistent with neuronal stress and injury occurring adjacent to sites of vascular compromise and neuroinflammation (**Figure 5c,d).**

**Figure 5:**
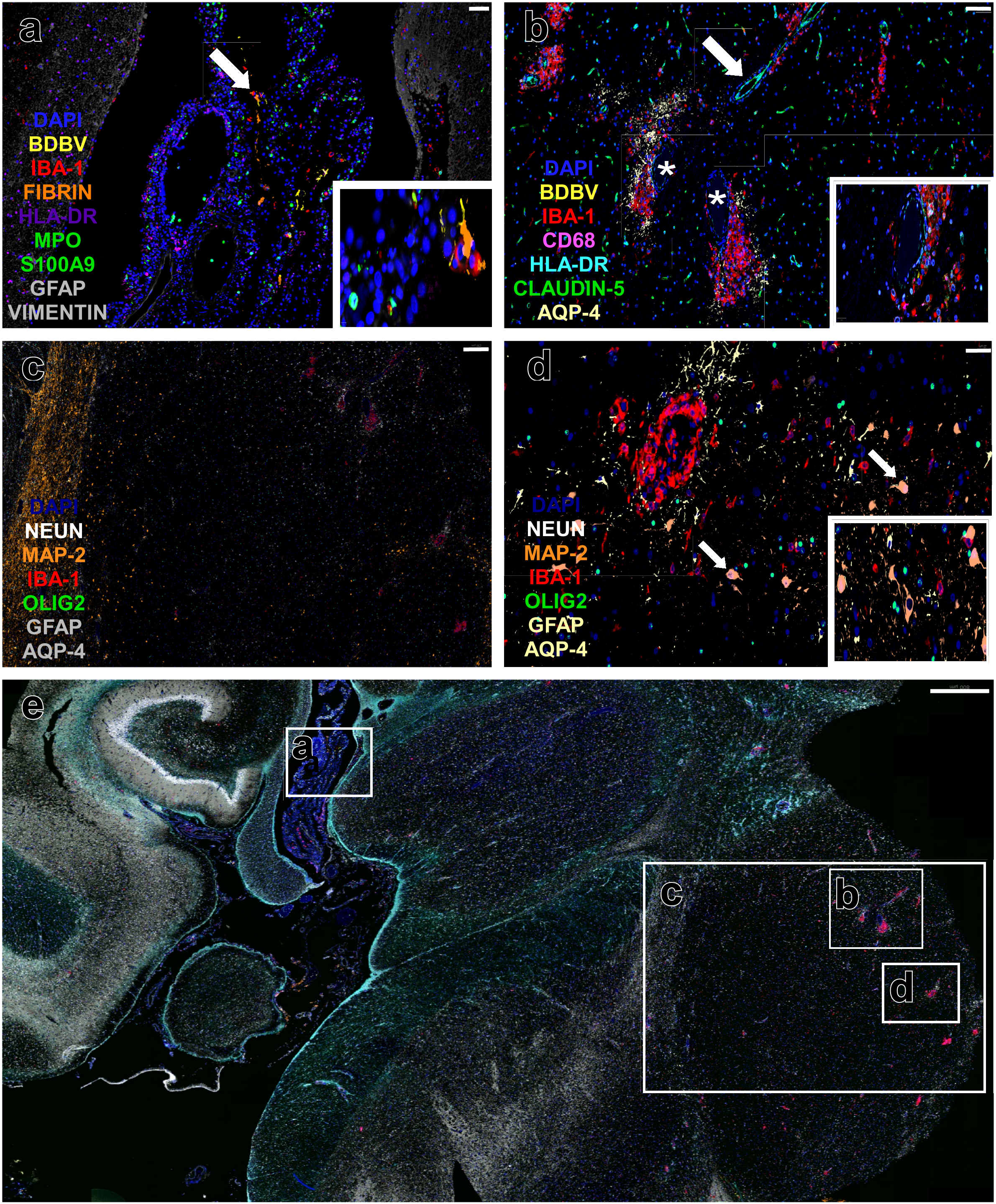
Multiplex immunofluorescence (mIF) of neuronal tissues of BDBV-infected NHP (CYNO2-6). **A)** Choroid plexus demonstrating abundant BDBV antigen-positive (yellow) cells localized within IBA-1+ macrophages (red), including activated IBA-1+/HLA-DR+ (purple) macrophages. Additional BDBV-positive cells are present within the choroid plexus epithelium (ependymal cells), identified by the absence of leukocyte, endothelial, and other structural markers. Moderate numbers of MPO+/S100A9+ (green) neutrophils and/or myeloid-derived suppressor cells (MDSCs) are present within vascular lumina and the choroid plexus stroma. Scant fibrin deposition (orange) is observed within the stroma. DAPI (blue) labels nuclei, GFAP (gray) identifies astrocytes within adjacent neuroparenchyma, and vimentin (gray) labels endothelial and stromal cells. Notably, vimentin expression is absent in the choroid plexus epithelium, likely reflecting downregulation associated with the marked inflammatory response. These findings demonstrate substantial viral antigen accumulation and inflammation within the choroid plexus, a key interface between the peripheral circulation and central nervous system. Scale bar = 50 μm. Inset higher magnification of BDBV+IBA-1+FIBRIN+ cells. **B)** Thalamus with multifocal perivascular inflammation characterized by rare BDBV-positive (yellow) IBA-1+ (red) macrophages/microglia and activated IBA-1+/HLA-DR+/CD68+ inflammatory cells (light blue and pink). Vessels associated with minimal inflammation exhibit strong Claudin-5 expression (white arrow), consistent with intact endothelial tight junctions and preservation of BBB integrity. In contrast, vessels surrounded by dense inflammatory infiltrates show markedly reduced or absent Claudin-5 labeling and are associated with increased Aquaporin-4 (AQP-4; light yellow) expression in astrocytic end feet (*), indicative of vascular leakage, edema, and BBB disruption. BDBV-positive macrophages are concentrated around these compromised vessels, suggesting viral-associated neuroinflammation contributes to loss of BBB integrity. Endothelial cells retaining strong Claudin-5 expression also exhibit increased HLA-DR labeling, consistent with endothelial immune activation. Inset higher magnification of inflamed vessel with perivascular BDBV+IBA-1+, CD68+, HLA-DR+ cells surrounding vessel with diminished claudin-5 and HLA-DR expression **C)** Thalamic regions containing perivascular inflammation demonstrate reduced neuronal density with sparse MAP2+ neurons, all lacking nuclear NeuN expression, consistent with neuronal stress, dysfunction, or degeneration. **D)** Higher magnification of inflamed thalamic regions highlighting MAP2+ neurons lacking NeuN nuclear staining (white arrows), further supporting neuronal injury associated with adjacent vascular and inflammatory lesions. Inset higher magnification of MAP+ NeuN-cells. **E)** Low-magnification overview of the temporal lobe demonstrating the spatial relationship between viral antigen, inflammatory infiltrates, vascular alterations, and neuronal populations. Insets correspond to panels A–D. Scale bar = 800 μm.

Collectively, BDBV-associated lesions were systemic and involved multiple organ systems, with the most severe pathology occurring in lymphoid tissues, MALT, and liver. Viral antigen and associated lesions were additionally detected within immune-privileged sites, including the brain, eye, and reproductive tract, most prominently in animals with prolonged disease courses, consistent with progressive viral dissemination during late-stage infection. Additional histopathologic descriptions and representative lesion images are provided in the **Supplementary Data** and **Figure 4g,h,k,l,o,p,q,r,s**.

### Transcriptomics and proteomics

To more thoroughly characterize the host response to BDBV infection, we performed targeted transcriptomics and proteomics on circulating blood and sera from CM enrolled in the natural history study (CYNO2-1 through −12). Naïve dimension reduction identified distinct disease stages following BDBV exposure: pre-response, early, middle, late, and recovery (**Supplementary Figure 6**). No significantly differentially expressed (DE) mRNAs or proteins were identified in pre-response samples (**Figure 6; Supplementary Figure**). In early-stage disease, we observed upregulation of interferon-stimulated gene (ISG) transcripts in CM regardless of outcome as well as increased CXCL10/IP-10 transcripts; no changes in proteins reached statistical significance (p_adj_ ≤ 0.05 and log_2_ fold-change > 1 or < −1).

**Figure 6:**
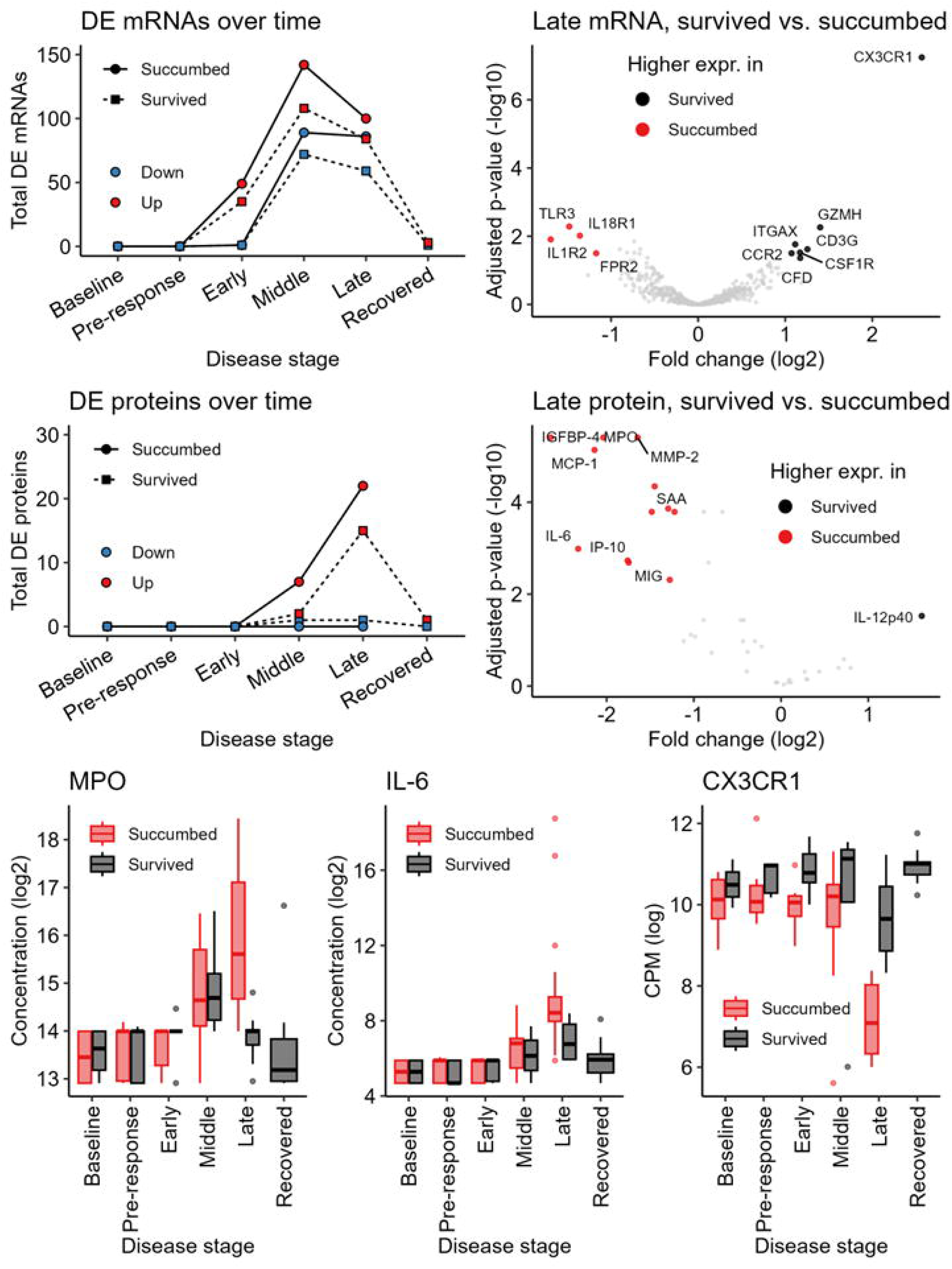
Transcriptomic and proteomic host response in NHP challenged with BDBV. Total significantly upregulated (red) and downregulated (blue) mRNA transcripts **(a)** or proteins **(b)** at each disease stage compared to pre-exposure baseline in CM that succumbed (circles) or survived (squares). Volcano plots of significantly upregulated mRNAs **(c)** or proteins **(d)** in survivor (black) or succumbed (red) CM during the late disease stage. Significantly DE transcripts had FDR-adjusted p-value < 0.05 and log_2_ fold change >1 or <−1. Box plots of log_2_-transformed concentrations (pg/mL) of MPO **(e)** and IL-6 **(f)** in succumbed (red) and survivor (black) CMs across disease stages.

Total DE mRNAs peaked in mid-disease. More ISGs were upregulated, as were cytokine mRNAs. Corresponding proteins increased, including CXCL10/IP-10, MCP-1, MRP8/14 (aka calprotectin or S100A8/9 complex), and tPA. Host response patterns began to diverge in late disease (**Figure 6c-d**). The proinflammatory response in succumbing animals continued to escalate, with additional acute-phase and neutrophil-associated mRNAs and proteins upregulated relative to pre-exposure baseline. Several of these analytes, including MPO and IL-6 (**Figure 6e-f**), were also significantly elevated relative to late-stage disease samples from surviving CM. In contrast, survivors had higher expression of transcripts associated with immune-mediated viral clearance (i.e., CX3CR1, GZMH, and CD3G). Differences between outcomes were only notable in late-stage disease; succumbing animals were dominated by the same S100A9/MPO signature observed in tissues. Survivors pivoted from the pro-inflammatory phenotype seen in mid-stage disease and expressed mRNAs and proteins consistent with viral clearance and immune regulation. Overall, aberrant myeloid-associated immune signatures drove a hyperinflammatory state contributing to the observed histopathology, uncontrolled viral replication, and fatal survival outcomes.

## Discussion

As filovirus outbreaks occur sporadically in resource poor settings conducting clinical trials of MCM can be challenging. The US Food and Drug Administration (FDA) Animal Rule offers a pathway for licensure of MCM where human efficacy studies are infeasible or unethical^38^.

However, only MCM for anthrax pre- and postexposure prophylaxis have been approved using the Animal Rule^39^. The ERVEBO vaccine for EBOV and the human monoclonal antibodies Inmazeb and Ebanga, also for EBOV, were all approved for human use based on clinical trials conducted in 2015 during the West African epidemic and in 2018 during the outbreak in the DRC, respectively^8^. These clinical trials were informed and benefited from preclinical work done in NHP models of Ebola virus disease (EVD)^40–44^. Given the magnitude of the current BDBV outbreak in DRC and Uganda it is likely that MCM being deployed will follow a similar path to licensure. Past and current studies in NHP models of BVD hopefully have helped and will continue to assist in the use of MCM during this outbreak and development of new MCM.

High levels of circulating BDBV RNA and infectious virus were detected in all 12 macaques in the natural history study. There was no association between the quantity of circulating BDBV RNA and survival, consistent with our previous work in BDBV-infected rhesus macaques^32^. There have been no reports during the few human BDBV outbreaks associating circulating viral load with patient outcome. In contrast to these BDBV NHP studies, high circulating levels of EBOV and SUDV RNA have been associated with poor outcome and lower levels with survival in humans and NHP^16,45–49^. Here, we also observed an extended disease course for BDBV in CM versus EBOV and SUDV consistent with previous studies and the 2012 BDBV outbreak. The slower disease course provides the host with more time to mount a protective adaptive immune response which may contribute to the lower case fatality rates noted for BDBV and also provides more time to administer therapeutic interventions. However, an extended disease course also means that infected individuals are likely able to transmit the virus to others for a longer period of time potentially making it harder to control outbreaks.

Collectively, findings from the temporal euthanasia study show a progression of events from initial infection of mononuclear phagocytic cells within lymphoid and vascular compartments at 4 DPI to widespread systemic dissemination by 8 DPI. IHC and spatial proteomics identified CD209+, CD68+, and/or HLA-DR+ macrophages and dendritic cells as early targets of BDBV consistent with previous findings for Ebola virus-infected CM^34^. These infected cells frequently colocalized with fibrin and infiltrating MPO+ neutrophils and S100A9+ myeloid-derived suppressor cells, consistent with the development of an active inflammatory response and early coagulopathy. BDBV antigen was localized not only to highly vascularized organs but also to multiple mucosal surfaces, exocrine glands, and epithelial compartments associated with potential viral shedding, including the conjunctiva, salivary glands, gastrointestinal tract, urinary bladder, reproductive tract, mammary tissues, and sebaceous glands. Concurrently, infection of immune-privileged tissues, including the eye, central nervous system, and reproductive organs, suggests that acute hematogenous dissemination may provide a mechanism for viral access to anatomical sites characterized by specialized immune regulation.

The natural history study further advanced findings from the temporal euthanasia study and showed that lesion severity and BDBV antigen burden generally peaked between 9 and 12 DPI, coinciding with extensive involvement of major target organs and systemic spread of infection. BDBV antigen was consistently identified within multiple epithelial and mucosal surfaces, including the oral cavity, tonsil, gastrointestinal tract, conjunctiva, nasal mucosa, urinary tract, skin, mammary gland, and reproductive tract, further supporting these tissues as potential sources of viral shedding and transmission during acute disease. Both surviving and succumbing CM exhibited the cytokine-driven proinflammatory immune response characteristic of EVD^50^ in early and mid-disease, with outcome-dependent differences only noted in late disease. While succumbing NHP only increased expression of neutrophilic and acute-phase markers until meeting endpoint criteria, survivors transitioned to a more controlled and targeted transcriptomic and proteomic profile in late disease, then returned to a near-baseline immune state by the study endpoint. Notably, BDBV antigen persisted in immune-privileged and sanctuary sites, including the brain, eye, and reproductive organs, particularly in animals surviving to later stages of disease. These findings mirror sites associated with viral persistence and long-term sequelae in human EBOV and SUDV survivors^51–54^ and suggest that infection of these tissues during acute disease may contribute to the development of post-Ebola syndrome, including neurologic, ocular, and reproductive complications observed following recovery. Persistence of EBOV has previously been noted in immune privileged tissues of NHP that had delayed disease courses or survived EBOV or SUDV infection^55–58^.

In summary, observations from these studies establish a histologic framework for understanding both potential routes of BDBV transmission and the early tissue events that may contribute to the development of persistent tissue reservoirs following acute infection. These data should be helpful in identifying potential targets for the development of new interventions and/or the optimization of MCM strategies currently in preclinical studies.

## Methods

### Challenge virus

BDBV isolate 200706291 (BDBV/But-811250 variant GenBank Accession ref: PZ485171) was obtained from a fatal human case in western Uganda during the 2007 outbreak^6^. The challenge stock used in this study was propagated on Vero E6 cells twice (passage 2 virus) and certified free of endotoxin and mycoplasma contamination; negative stain transmission electron microscopy confirmed that virus particles were consistent in morphology with an orthoebolavirus.

### Nonhuman primate challenge

A total of 21 healthy, research-naïve, captive bred cynomolgus macaques (*Macaca fascicularis*) ∼ 3.9-7.5 years of age and weighing ∼ 2.9-6.9 kg were obtained from a commercial vendor (PreLabs, Lehigh Acres, FL). Studies were performed to 1) assess the progression of events in tissues during BDBV infection and 2) to assess the natural history of BDBV infection. Efforts were made to balance the sex distribution in each cohort (the sex of each individual animal is provided in **Supplementary Tables 1 & 2**). All 21 macaques were exposed by intramuscular (i.m.) injection in the left quadriceps to 1000 PFU of BDBV (200706291 isolate). For the temporal pathogenesis work three animals per day were euthanized on days 2, 4, and 8 after exposure to BDBV to collect blood and tissues. For the natural history work 12 animals were allowed to progress to the predetermined study endpoint of 28 DPI or until they met criteria for euthanasia.

All animal work was approved by the UTMB Institutional Animal Care and Use Committee (IACUC, approval number D16-00202). All macaques were monitored daily and scored for disease progression with an internal BDBV humane endpoint scoring sheet approved by the UTMB IACUC. UTMB facilities used in this work are accredited by the Association for Assessment and Accreditation of Laboratory Animal Care International and adhere to principles specified in the eighth edition of the Guide for the Care and Use of Laboratory Animals, National Research Council. The scoring changes measured from baseline included posture and activity level, attitude and behavior, food intake, respiration, and disease manifestations, such as visible rash, hemorrhage, ecchymosis, or flushed skin. A score of ≥ 9 indicated that an animal met the criteria for euthanasia.

### Hematology and serum biochemistry

Total white blood cell counts, white blood cell differentials, red blood cell counts, platelet counts, hematocrit values, total hemoglobin concentrations, mean cell volumes, mean corpuscular volumes, and mean corpuscular hemoglobin concentrations were analyzed from blood collected in tubes containing EDTA using a laser based hematologic analyzer (Beckman Coulter). Serum samples were tested for concentrations of albumin, amylase, alanine aminotransferase (ALT), aspartate aminotransferase (AST), alkaline phosphatase (ALP), blood urea nitrogen (BUN), calcium, creatinine (CRE), C-reactive protein (CRP), gamma-glutamyltransferase (GGT), glucose, total protein, and uric acid by using a Piccolo point-of-care analyzer and Biochemistry Panel Plus analyzer discs (Abaxis).

### RNA isolation

On the specified procedure days, blood was collected from each macaque by femoral venipuncture into EDTA vacutainer tubes (BD Biosciences, San Jose, CA). An aliquot of EDTA-treated whole blood (100 μl) was diluted with 600 μl of AVL inactivation buffer (Qiagen, Hilden, Germany), and RNA was extracted using a Viral RNA mini-kit (Qiagen) according to the manufacturer’s instructions.

### Viral Load Determination

One-Step Probe RT-qPCR kits (Qiagen) and CFX96 system/software (BioRad) were used to determine BDBV viral copies. To detect viral RNA, we targeted the BDBV VP35-intergenic region with primer pairs and a 6-carboxyfluorescein (6FAM)-5′ CGCAACCTCCACAGTCGCCT 3′-6 carboxytetramethylrhodamine (TAMRA). Thermocycler run settings were 50°C for 10 minutes; 95°C for 10 seconds; and 40 cycles of 95°C for 10 seconds plus 57°C for 30 seconds. Integrated DNA Technologies synthesized all primers and Life Technologies customized probes. Representative BDBV genomes were calculated using a genome equivalent standard, which takes into account Avogadro’s number and the molecular weight of the BDBV genome. The limit of detection for this assay is 1000 copies/ml.

Virus titration was performed for BDBV by plaque assay with Vero E6 cells (ATCC, CRL-1586) from all plasma samples as previously described^23,24^. Briefly, increasing 10-fold dilutions of the samples were adsorbed to Vero E6 monolayers in duplicate wells (200 µL) and overlaid with 0.8% agarose in 1× Eagle’s minimum essentials medium (EMEM) with 5% fetal bovine serum and 1% penicillin-streptomycin. After a 7-day incubation at 37°C/5% CO2, neutral red stain was added, and plaques were counted after a 24-48-hour incubation. The limit of detection for this assay was 25 PFU/mL.

### IgG ELISA

Sera collected at the indicated time points were tested for BDBV GP-specific IgG antibodies by ELISA. MaxiSorp 96-well plates (44204 ThermoFisher, Rochester, NY) were coated overnight with 0.08uL/mL of recombinant BDBV GPΔTM (ΔTM: transmembrane region absent; Integrated Biotherapeutics, Gaithersburg, MD) in a sodium carbonate/bicarbonate solution (pH 9.6). Antigen-adsorbed wells were subsequently blocked with 4% bovine serum antigen (BSA) in 1 x PBS for at least two hours. Sera were initially diluted 1:100 and then two-fold through 1:25600 in ELISA diluent (1% BSA in 1× PBS, and 0.2% Tween-20). After a one-hour incubation, cells were washed five times with wash buffer (1 x PBS with 0.2% Tween-20) and incubated for an hour with a 1:15000 dilution of horseradish peroxidase (HRP)-conjugated anti-monkey IgG antibody (BioSynth, Gardner, MA). O-phenylenediamine (OPD) Substrate tablet (Thermo Scientific; 34006) dissolved in Stable Peroxide Substrate Buffer (Thermo Scientific; 34062) was added to the wells after five additional washes to develop the colorimetric reaction. The reaction was stopped with 2.5M sulfuric acid 10 minutes after OPD addition and absorbance values were measured at a wavelength of 492 nm on a Cytation 5 (Agilent BioTek, Santa Clara, CA). Absorbance values were normalized by subtracting uncoated from antigen-coated wells at the corresponding serum dilution. End-point titers were defined as the reciprocal of the last adjusted serum dilution with a value ≥ 0.15.

### Histopathology and immunohistochemistry

Necropsy was performed on all subjects in the BSL-4. Tissue samples for histopathologic and immunohistochemical (IHC) examination were immersed in 10% neutral buffered formalin for at least 21 days followed by a change of formalin before removal from the BSL-4. Inactivated tissue samples were processed in a BSL-1. Tissue sections were deparaffinized and rehydrated through xylene and graded ethanols. Slides went through heat antigen retrieval in a steamer at 95°C for 20 minutes in Sigma Citrate Buffer, pH6.0, 10x (Sigma Aldrich, St. Louis, MO). The tissue sections were processed for IHC using the Thermo Autostainer 360 (ThermoFisher, Kalamazoo, MI). Specific anti-BDBV immunoreactivity was detected using an anti-BDBV GP primary antibody at a 1:2000 dilution for 60 min. Secondary antibody used was biotinylated goat anti-rabbit IgG (Vector Laboratories, Burlingame, CA #BA-1000) at 1:200 for 30 min followed by Vector Streptavidin Alkaline Phosphatase at a dilution of 1:200 for 15 minutes (Vector Laboratories #SA-5100). Slides were developed with ImmPact Red Substrate Kit (Vector Laboratories #SK-5105) for 20 minutes and counterstained with hematoxylin for 30 seconds.

### Multiplex immunofluorescence (mIF)

mIF was performed using the PhenoCycler-Fusion 2.0 platform (Akoya Biosciences/Quanterix) according to the manufacturer’s instructions. The assay was optimized and validated prior to use and included markers for: BDBV, IAB-1, Fibrin, HLA-DR, MPO, S100A9, GFAP, CD68, CD1d, Claudin-5, AQP4, NeuN, MAP-2, Olig2, Vimentin, CD209, and BCL-2. Detailed optimization parameters for antibodies are outlined in **Supplementary Table 5**. Appropriate positive and negative controls are included in each staining run to verify antibody specificity and staining performance. All tissues were evaluated by a board-certified veterinary pathologist using consistent criteria to support reproducibility and transparent reporting of pathological findings.

### Targeted transcriptomics

Whole blood RNA was used to quantify 770 immune-associated mRNAs via the Nanostring nCounter NHP Immunology v2 panel (Bruker #115000276). mRNAs were quantified with the nCounter SPRINT before importing into nSolver v4.0, and a background-thresholded count matrix was generated using the default parameters. Differential expression analyses were performed with limma v3.66.0^59^ in R v4.5.3^60^. P-values <0.05 were considered significant, and mRNAs were subject to an additional threshold of log_2_ fold change >1 or <−1.

### Circulating proteomics

Circulating chemokines, cytokines, and other protein markers associated with inflammation were measured via LEGENDplex™ bead-based multiplex immunoassay panels obtained from BioLegend. Gamma-irradiated serum samples were assessed in duplicate for NHP Inflammation V02 (#741491, 1:4 dilution), Human Fibrinolysis (#740761, 1:40,000 dilution), Human Vascular Inflammation (#740590, 1:1000 dilution), Human Thrombosis (#740892, 1:50 dilution), and NHP Chemokine/Cytokine (#740388, 1:4 dilution) panels according to manufacturer instructions. Assay standards were prepared in batches and aliquoted across all plates to ensure batch-to-batch consistency. All optional wash steps were incorporated into the workflow to reduce background signal. Assay samples were analyzed on an Accuri C6 Plus flow cytometer (BD Biosciences). The raw .fcs files from each assay plate were imported into LEGENDplex Qognit cloud-based Data Analysis Software Suite (BioLegend), which determined analyte concentrations in experimental samples via 5-parameter logistic regression curve fitting to each assay standard curve. Analyte concentration data were exported from Qognit and analyzed in R using limma v3.66.0^59^.

### Statistical analysis

Details about the specific statistical methods used for each comparison are listed in the main text, individual figure legends, and/or the relevant Methods subsection. For data generated by Nanostring targeted transcriptome profiling and LEGENDplex bead-based multiplex assays, statistical analyses were performed using limma v3.66.0^59^ in R v4.6.0^60^; analysis code is available on GitHub (https://github.com/geisbert-lab/bdbv2007-serial-sac). Analyses were performed using GraphPad Prism v11.0.2 and Microsoft Excel.

## Supporting information

Sipplementary Data

## Data sharing and availability

The genome sequence of the BDBV challenge stock is available in GenBank (PZ485171). Targeted transcriptomic data are deposited in GEO (Accession number TBD). All other data are presented in the Article and Appendix.

## Author contributions

RWC and TWG conceived and designed the experiment. CFB and TWG secured the funding. JBG and DJD performed the challenges. ANP, JBG, DJD, RWC, and TWG performed procedures and conducted clinical observations. KNA and VB performed clinical pathology assays. KNA performed the PCR assays. JBG performed the BDBV plaque assays. RO performed the ELISAs. JT and CW performed the gene expression assays and JT performed the transcriptomic analyses. DDP performed the bead-based multiplex assays and proteomic analysis. NSD and AL performed the IHC and mIF assays. KAF performed gross pathologic, histologic, immunohistochemical, and spatial protemomics analyses of the data. All authors analyzed the data. TWG wrote the paper with additions from DDP, JT, ANP, and KAF. DDP, JT, ANP, and KAF prepared the Figures and ANP prepared the Tables. CFB and RWC edited the manuscript. All authors had access to the data and approved the final version of the manuscript.

## Acknowledgments

This study was supported by the Department of Health and Human Services, National Institutes of Health grants U19AI109945 to CFB and U19AI142785 to TWG, UTMB Department of Microbiology and Immunology funds to TWG, and UC7AI094660 for BSL-4 operations support of the Galveston National Laboratory. The authors wish to thank the UTMB Animal Resource Center for husbandry support of laboratory animals. Opinions, interpretations, conclusions, and recommendations are those of the authors and are not necessarily endorsed by the University of Texas Medical Branch.

