## Supplementary material for "Pathogenesis and natural history of the Bundibugyo species of *Orthoebolavirus* in nonhuman primates": Sipplementary Data

**Supplementary Data**

**Gross lesions and histopathology**

Pulmonary lesions consisted of mild to moderate interstitial pneumonia through 15 DPI, characterized by expansion of alveolar septa by inflammatory leukocytes and increased alveolar macrophages, a subset of which contained viral antigen (**Figure 4g**). Pulmonary lesions and immunolabeling were absent in animals surviving beyond 15 DPI, including all survivors. Seven of eight succumbed animals demonstrated viral antigen within endocardial and myocardial interstitial cells in the absence of significant inflammation. No cardiorespiratory lesions or BDBV antigen were observed in surviving animals (**Supplementary Figure 5f**).

Renal lesions in succumbed animals were mild to moderate and consisted of interstitial nephritis with degeneration of proximal convoluted tubule epithelium. BDBV antigen was present within interstitial inflammatory cells, glomerular leukocytes, vascular endothelium, and the subepithelial stroma of the renal pelvis extending into the ureter (**Figure 4k**). In the urinary bladder, antigen-positive inflammatory cells and transitional epithelial cells were present in all but two succumbed animals. Extensive dissemination of viral antigen was observed throughout the reproductive tract. In females, BDBV-positive mononuclear cells were present within the theca interna and externa of follicles at multiple stages of development (**Figure 4r**), while the uterus contained dense bands of BDBV-positive stromal cells immediately beneath the glandular epithelium (**Figure 4s**). In males, BDBV-positive mononuclear cells were identified within the vascular interstitium of the testes and epididymides and within inflammatory nodules of the prostate (**Figure 4q**). Endocrine tissues were widely affected and exhibited varying degrees of degeneration, hemorrhage, or necrosis accompanied by viral antigen within pancreatic islets, thyroid follicular and parafollicular cells, parathyroid tissues, adrenal cortical cells (**Figure 4h**) and infiltrating leukocytes, and cells of the anterior and posterior pituitary glands (**Figure 6t**). Two surviving animals exhibited minimal perivascular to interstitial lymphohistiocytic infiltrates within the kidney without detectable BDBV antigen (**Supplementary Figure 5g,h,m,n**).

Ocular involvement ranged from minimal to moderate uveitis. Mild uveitis with antigen-positive mononuclear cells within the drainage angle and ciliary body was observed in CYNO2-4, CYNO2-5, and CYNO2-6. In CYNO2-10 and CYNO2-12, BDBV antigen was additionally present within pigmented and nonpigmented ciliary body epithelium (**Figure 4l**). More advanced ocular disease was present in CYNO2-7 and CYNO2-9, in which BDBV-positive inflammatory cells extended into the vitreous chamber and/or optic nerve. Beyond the eye, BDBV-positive inflammatory cells and epithelial cells were frequently identified within the conjunctiva, nasal mucosa, haired skin, and mammary gland. Cutaneous lesions were characterized by immunoreactive mononuclear cells within the dermis, hair follicles, vascular endothelium, sebaceous glands, and epidermis (**Figure 4o**). Mammary glands contained BDBV-positive inflammatory infiltrates surrounding and infiltrating ducts, and one animal exhibited intraductal accumulations of inflammatory and cellular debris (**Figure 4p**). No significant ocular, cutaneous, or mucosal lesions and no BDBV antigen labeling were observed in surviving animals (**Supplementary Figure 5j,l**).

BDBV infection in CM resulted in progressive multisystemic disease characterized by widespread viral dissemination, inflammation, degeneration, and necrosis involving lymphoid, gastrointestinal, hepatic, endocrine, neurologic, reproductive, ocular, and cutaneous tissues. Lesion severity and viral antigen burden generally peaked between 9 and 12 DPI, coinciding with extensive involvement of major target organs and systemic spread of infection. BDBV antigen was consistently identified within multiple epithelial and mucosal surfaces, including the oral cavity, tonsil, gastrointestinal tract, conjunctiva, nasal mucosa, urinary tract, skin, mammary gland, and reproductive tract, supporting these tissues as potential sources of viral shedding and transmission during acute disease. Notably, BDBV antigen persisted in immune-privileged and sanctuary sites, including the brain, eye, and reproductive organs, particularly in animals surviving to later stages of disease. These findings mirror sites associated with viral persistence and long-term sequelae in human Ebola virus disease survivors and suggest that infection of these tissues during acute disease may contribute to the development of post-Ebola syndrome, including neurologic, ocular, and reproductive complications observed following recovery.
