## Supplementary Figures for "Pathogenesis and natural history of the Bundibugyo species of *Orthoebolavirus* in nonhuman primates"

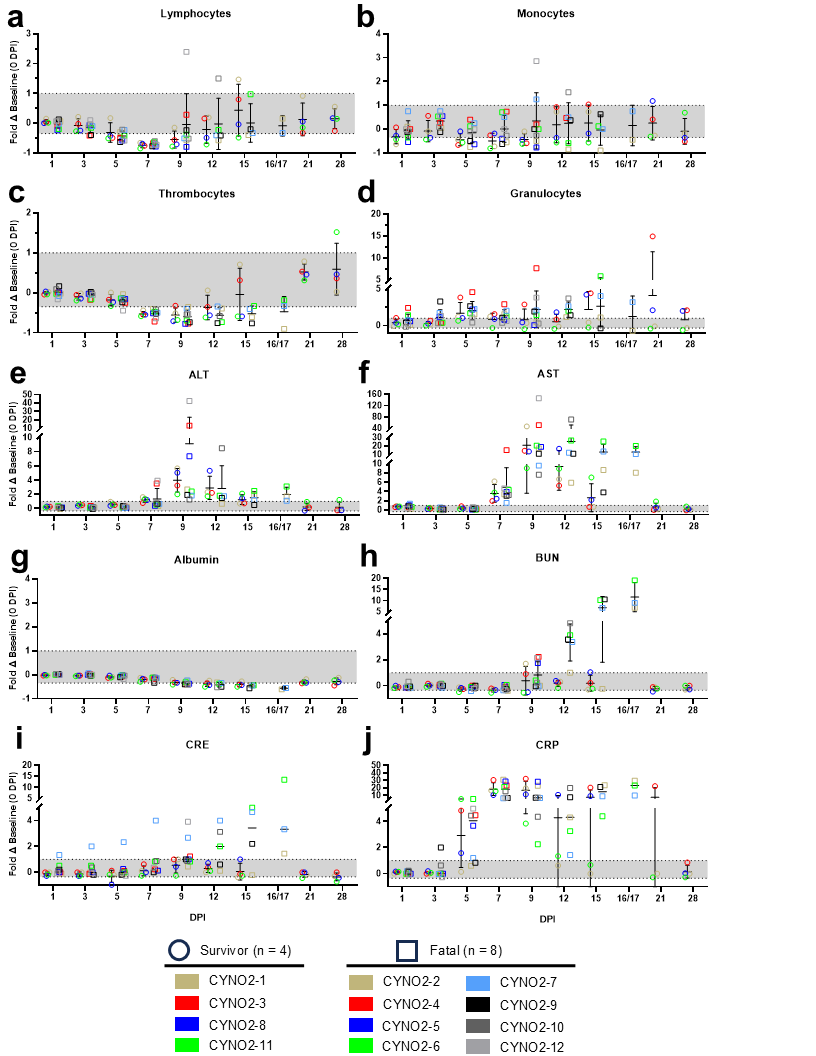

**Supplementary Figure 1: Selected hematological and serum biochemical markers from cynomolgus macaques challenged with BDBV.** Whole blood was obtained at pre-determined timepoints (0, 3, 5, 7, 9, 12, 15, 21, 28 DPI) or at the terminal timepoint from all macaques, and complete blood cell counts and serum biochemical analysis were performed. Individual data points depict the fold change at each time point compared to baseline (0 DPI). **(a)** lymphocytes; **(b)** monocytes; **(c)** thrombocytes (platelets); **(d)** granulocytes; **(e)** ALT; **(f)** AST; **(g)** albumin; **(h)** blood urea nitrogen (BUN); **(i)** creatinine (CRE); **(j)** c-reactive protein (CRP). For all panels, the middle line shows the mean, and the error bars depict ± SD. The area indicated in gray indicates the normal range of variation from baseline; fold change values ≤ -0.35 (-0.25 for ALT, AST, albumin, BUN, CRE, and CRP) or ≥ 1 are considered clinically significant. Raw cell counts and serum analyte measurements were obtained once per animal per timepoint. Statistical analysis was performed using independent Mann-Whitney U-tests and corrected for multiple comparisons using the Holm-Šídák method.

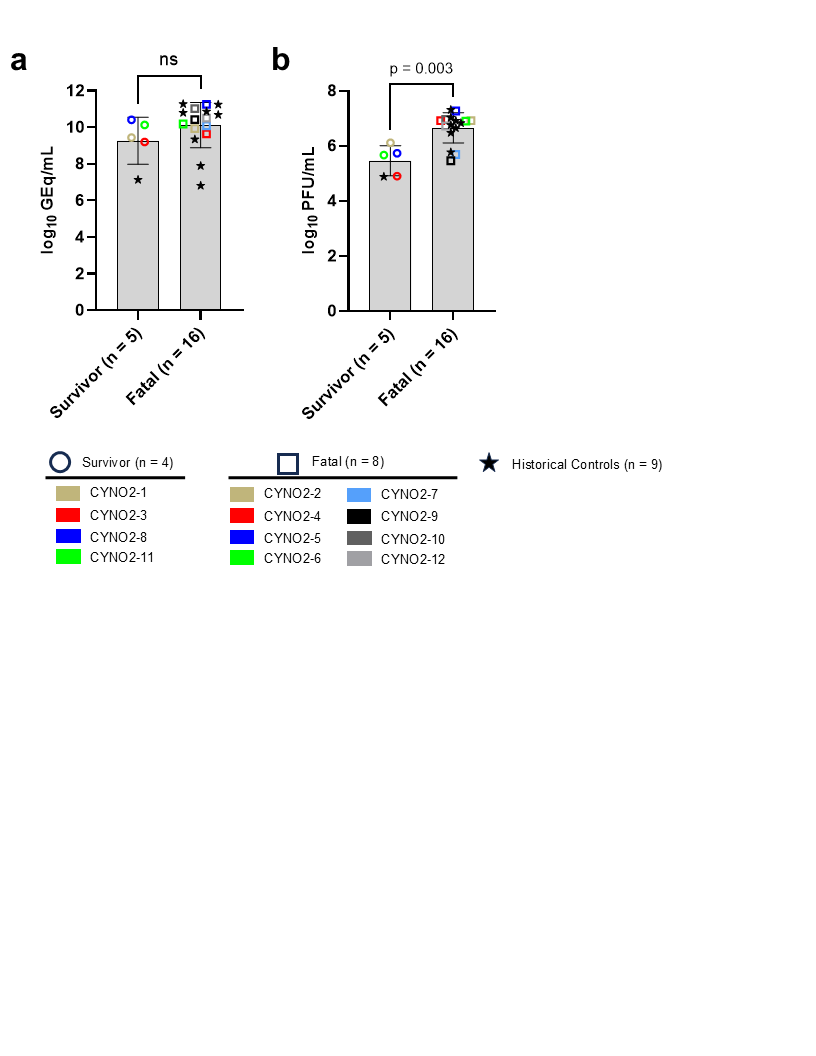

**Supplementary Figure 2: Comparison of peak circulating vRNA and infectious virus titers in macaques that survived BDBV challenge versus those that succumbed to lethal disease. (a)** Comparison of peak circulating vRNA abundance**; (b)** Comparison of peak circulating infectious virus titers. Plotted data points represent the mean of two replicate assays. Bars represent the geometric mean ± geometric SD. For both panels, data from historical positive control (HC) cynomolgus macaques (n = 9) was included for statistical comparison. Significance was tested using the non-parametric Mann-Whitney U-test. Reported p-value is two-tailed. ns = not significant.

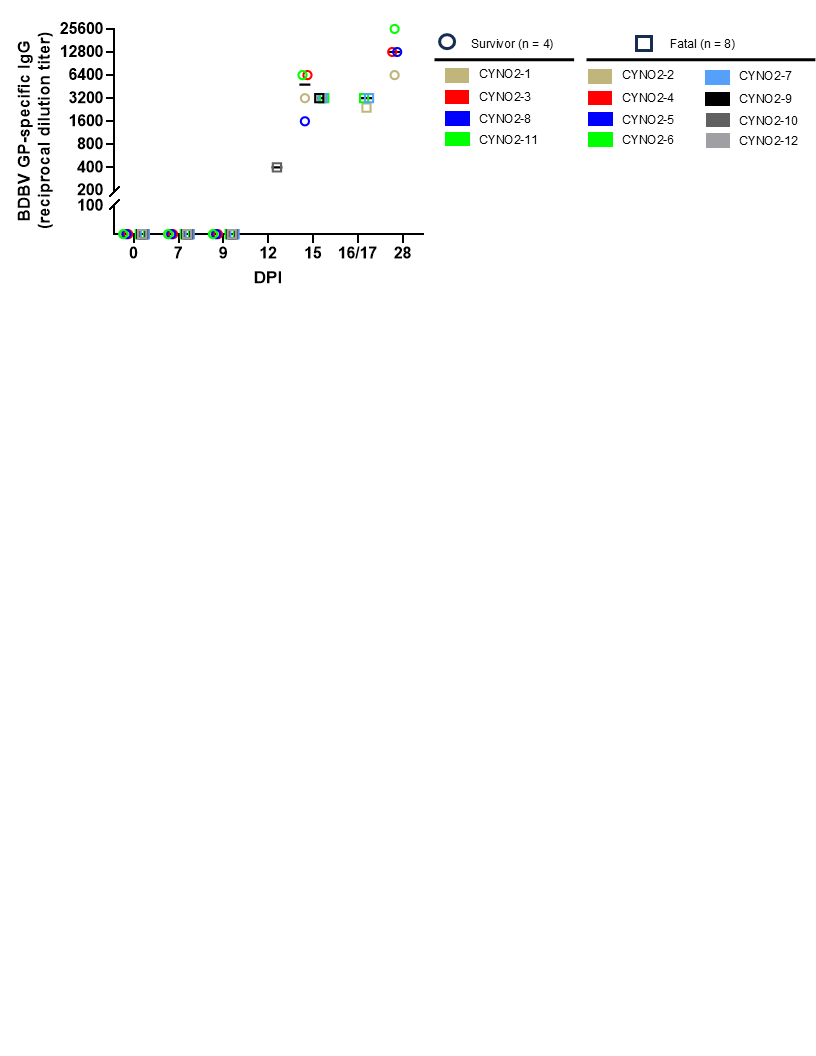

**Supplementary Figure 3: Humoral responses in BDBV-infected NHP**. Serum samples from were tested for circulating anti-BDBV GP-specific IgG by indirect ELISA. Line graphs depicting the average reciprocal dilution titer for individual subjects at each timepoint (0, 7, 9, 15, and 28 DPI) are shown. DPI, days post infection.

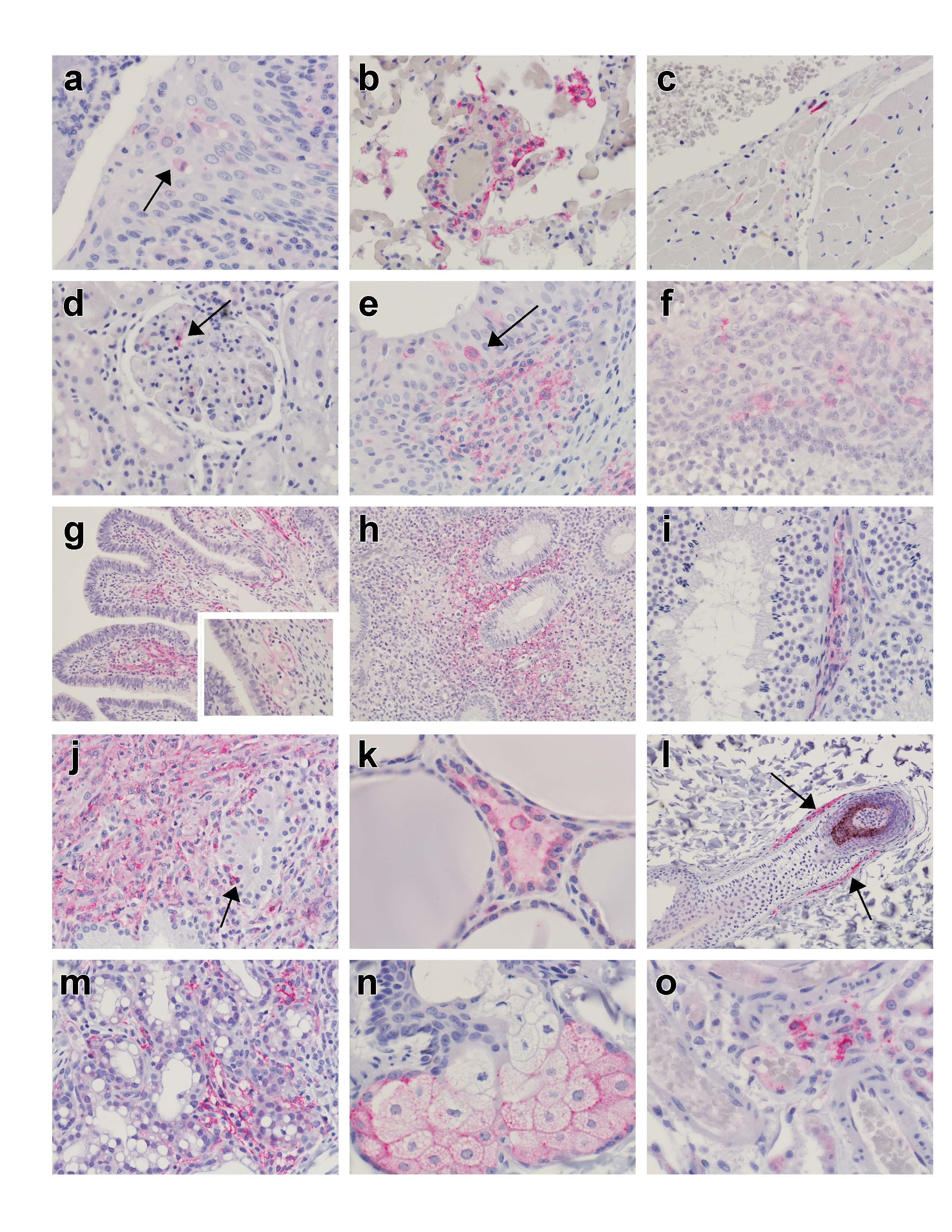

**Supplementary Figure 4: IHC of tissues from BDBV-infected NHP.** Representative IHC images for anti-BDBV GP antibody (red) in cynomolgus macaques (cyno) challenged with BDBV and euthanized on 8 DPI (**a-o**). Images captured at 20x magnification (**g,h,l**), 40x magnification (**b,d-f,i,j,m**), and 60x magnification (**a,g** inset, **k,n,o**).

a)	Tonsil, 60x, CYNO1-7: Scattered IHC positive cells within the subepithelial stroma and the epithelium (arrow).

b)	Lung, 40x, CYNO1-7: Minimal expansion of alveolar septa surrounding a small caliber vessels with mononuclear cells of which some are IHC positive.

c)	Heart, 40x, CYNO1-7: Scattered interstitial cells within the myocardium.

d)	Kidney, 40x, CYNO1-7: IHC positive mononuclear cells within the capillary bed of the glomerular tuff (arrow).

e)	Urinary Bladder, 40x, CYNO1-7: IHC positive mononuclear cells within the subepithelial stroma and the transitional epithelium (arrow).

f)	Ovary, 40x, CYNO1-8: IHC positive cells within the theca interna and externa.

g)	Infundibulum of the fallopian tube, 20x, CYNO1-8: Expansion of the stroma with Lymphohistiocytic inflammation with prominent IHC positivity. Inset, 60x-higher magnification with IHC positive endothelium.

h)	Uterus, 20x, CYNO1-8: Dense bands of IHC positive cells within the uterine stroma subjacent to the uterine gland epithelium

i)	Testis, 40x, CYNO1-7: IHC positive mononuclear interstitial cells of the testis

j)	Prostate, 40x, CYNO1-9: Dense bands of IHC positive cells within the prostatic stroma that abuts and transverses the glandular epithelium (arrow).

k)	Thyroid, 60x, CYNO1-7: IHC positive parafollicular and interstitial cells of the thyroid.

l)	Eyelid, 20x, CYNO1-7: IHC positive connective tissue sheaths surrounding hair follicles (arrows).

m)	Mammary gland/haired skin, 40x, CYNO1-9: IHC positive inflammatory infiltrates surrounding mammary ducts.

n) Eyelid, 60x, CYNO1-9: IHC positive sebaceous glands of the haired skin.

o)	Brain/Choroid Plexus, 60x, CYNO1-9: IHC positive cells within the choroid plexus of the ventricles.

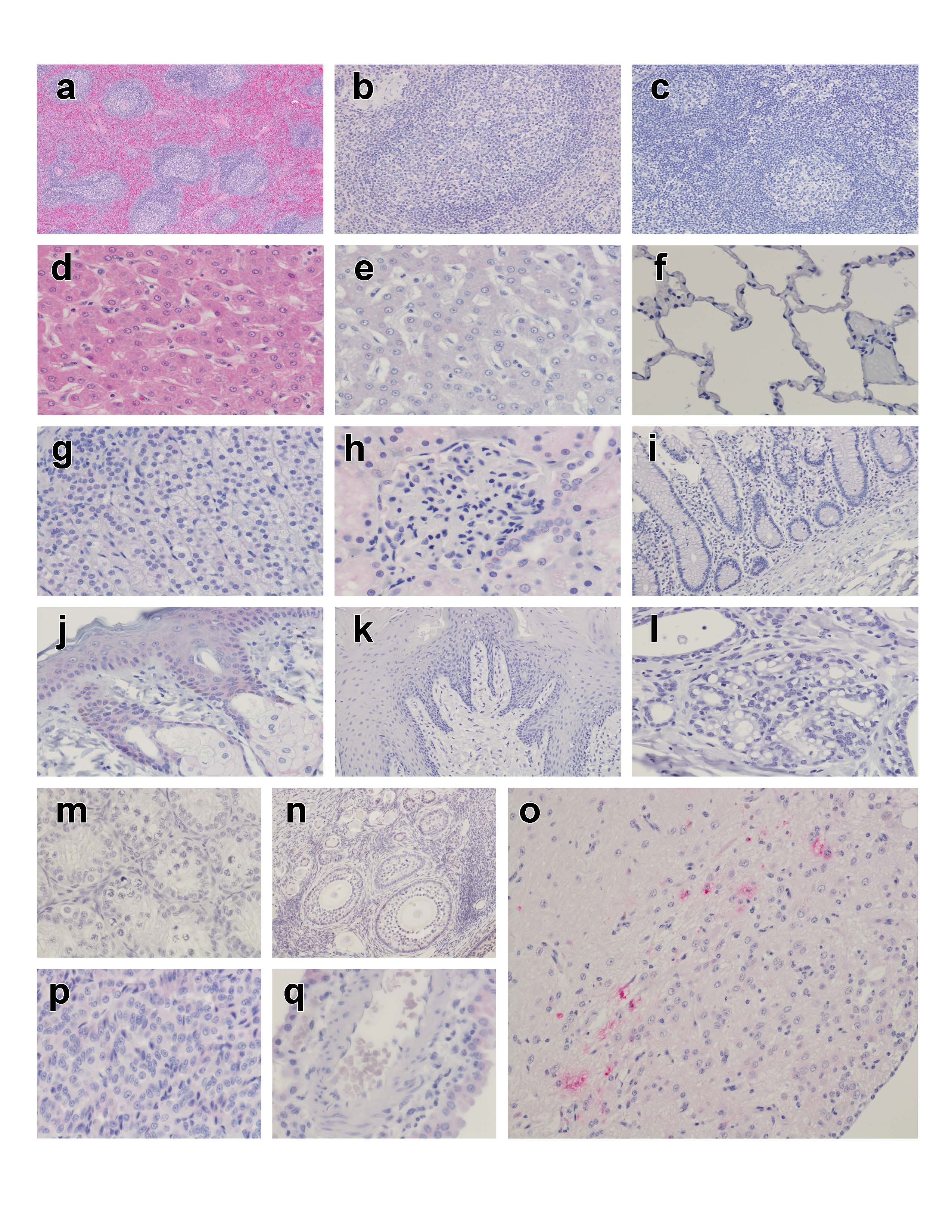

**Supplementary Figure 5: Histopathology and IHC of tissues from BDBV-infected NHP.** Representative H&E and IHC images for anti-BDBVGP antibody (red) in cynomolgus macaques (cyno) that survived challenge with BDBV. Images captured at 4x magnification (a), 20x magnification (b,c,j, k, n, o), 40x magnification (d,e,f,g,j,l,m), and 60x magnification (h,p,q).

a) Spleen, 4x, H&E, 130309, Day 28: No appreciable lesion.
b) Spleen, 20x, 130309, Day 28: No appreciable IHC labeling.
c) Lymph Node (Mesenteric), 20x, 130309, Day 28: No appreciable IHC labeling.
d) Liver, 40x, H&E, 130309, Day 28: No appreciable lesion.
e) Liver, 40x, 130309, Day 28: No appreciable IHC labeling.
f) Lung, 40x, 130309, Day 28: No appreciable IHC labeling.
g) Adrenal Gland, 40x, 130309, Day 28: No appreciable IHC labeling.
h) Kidney, 60x, 130309, Day 28: No appreciable IHC labeling.
i) Intestine, 20x, 130309, Day 28: No appreciable IHC labeling
j) Haired skin of the face, 130309, Day 28: No appreciable IHC labeling
k) Tongue, 20x, 14010906, Day 28: No appreciable IHC labeling
l) Mammary gland, 1311160, Day 28: No appreciable IHC labeling
m) Testis, 130309, Day 28: No appreciable IHC labeling
n) Ovary, 1401906, Day 28: No appreciable IHC labeling
o) Brainstem, 14010906, Day 28: Few scattered cells within the neuroparenchyma near the area postrema (red)
p) Brain, Pituitary gland, 130309, Day 28: No appreciable IHC labeling
q) Brain, Choroid Plexus, 1311160, Day 28: No appreciable IHC labeling

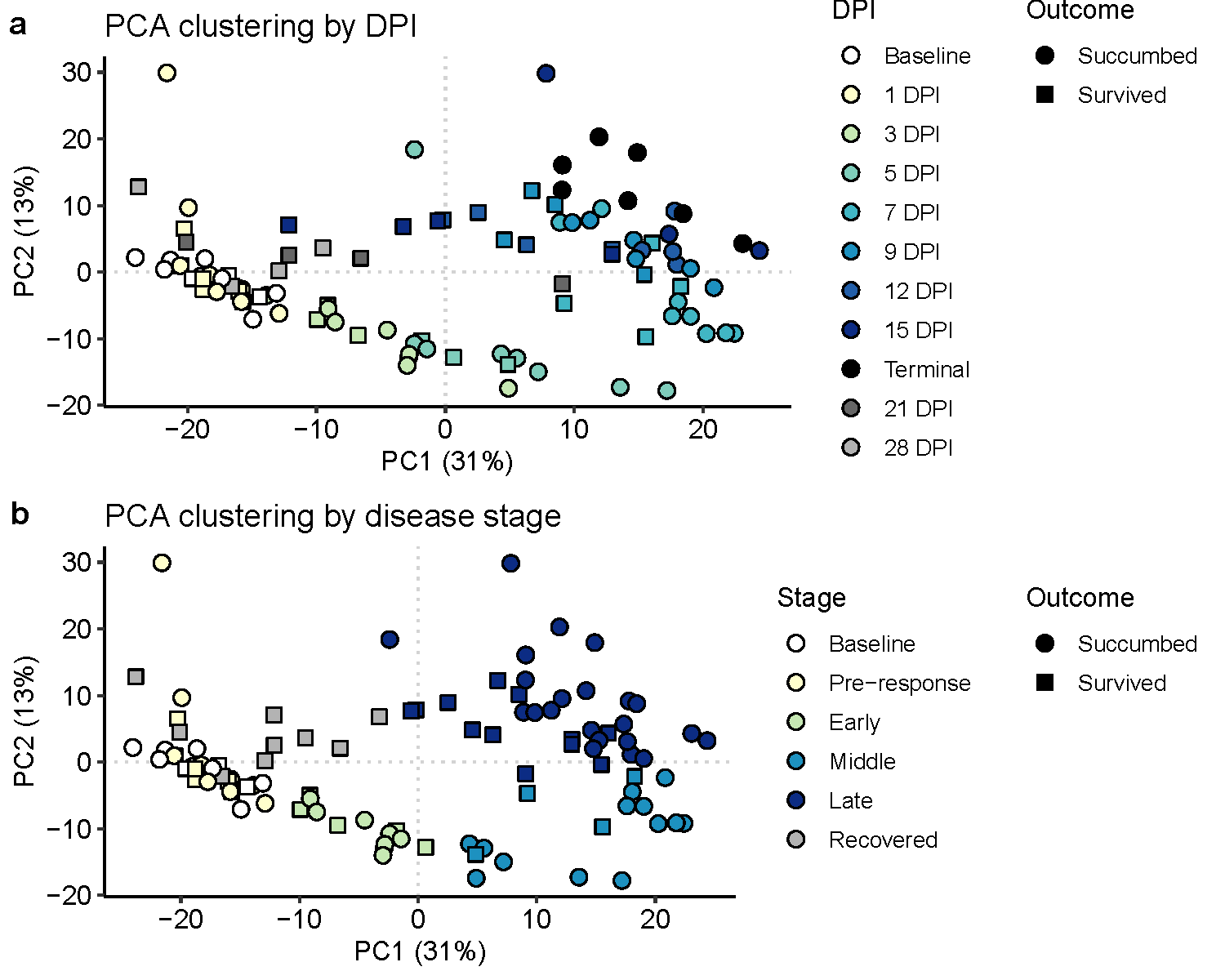
**Supplementary Figure 6:** Naïve clustering of BDBV-infected CMs (n=12) via PCA by DPI **(a)** or disease stage **(b)**.

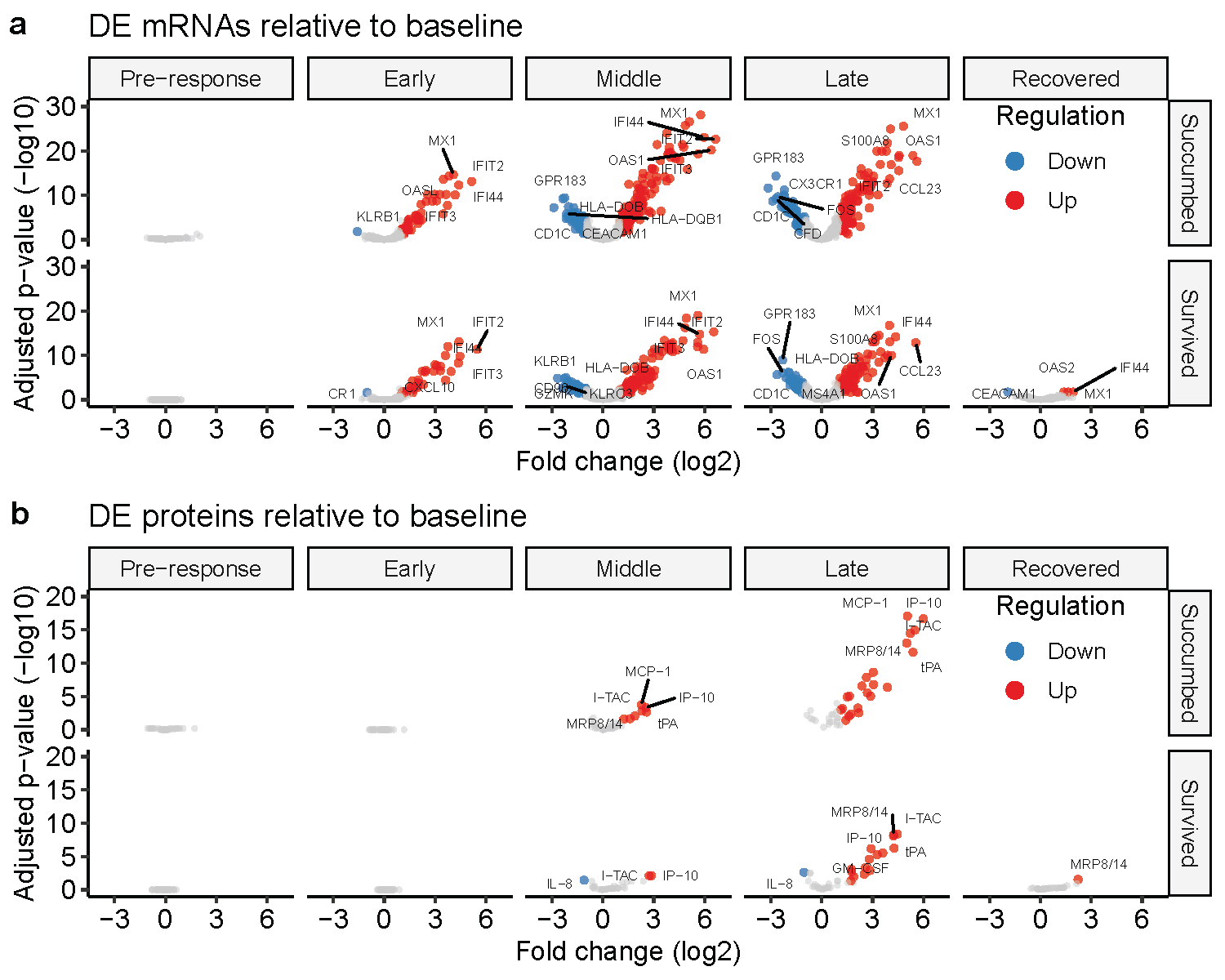

**Supplementary Figure 7:** Volcano plots of significantly differentially expressed mRNAs **(a)** or proteins **(b)** in circulation at distinct disease stages relative to pre-exposure baseline. For each disease stage, the log_2_ fold change was compared to baseline values of all n=12 CMs. Significantly DE transcripts had FDR-adjusted p-value < 0.05 and log_2_ fold change >1 or <-1.
