## Supplementary Tables for "Pathogenesis and natural history of the Bundibugyo species of *Orthoebolavirus* in nonhuman primates"

**Supplementary Table 1. Clinical description and outcome of cynomolgus macaques challenged with Bundibugyo virus (BDBV).**

| **Subject No.** | **Sex** | **Cohort** | **Clinical illness** | **Clinical pathology** |
| --- | --- | --- | --- | --- |
| CYNO1-1 | M | 2 DPI | None. Subject survived to study endpoint (d2). | AST ↑ (d1, 2). |
| CYNO1-2 | F | 2 DPI | None. Subject survived to study endpoint (d2). | Monocytosis (d1); granulocytosis (d1). |
| CYNO1-3 | F | 2 DPI | None. Subject survived to study endpoint (d2). | Monocytopenia (d2). |
| CYNO1-4 | M | 4 DPI | None. Subject survived to study endpoint (d4). | Monocytopenia (d2, 4); granulocytosis (d2, 4); ALT ↑ (d4). |
| CYNO1-5 | F | 4 DPI | None. Subject survived to study endpoint (d4). | Lymphocytopenia (d4); monocytopenia (d2, 4); granulocytosis (d4); CRP ↑ (d4). |
| CYNO1-6 | F | 4 DPI | None. Subject survived to study endpoint (d4). | Granulocytosis (d2, 4); |
| CYNO1-7 | M | 8 DPI | Decreased appetite (d5, 7); anorexia (d8); depression (d8). Subject survived to study endpoint (d8). | Lymphocytopenia (d8); thrombocytopenia (d8); hypoamylasemia (d8); ALT ↑ (d8); AST ↑ (d8); ALP ↑ (d8); CRP ↑↑↑ (d8). |
| CYNO1-8 | F | 8 DPI | Decreased appetite (d6); anorexia (d7, 8); depression (d7, 8); hunched posture (d7, 8). Subject survived to study endpoint (d8). | Lymphocytopenia (d8); thrombocytopenia (d8); granulocytosis (d4, 8); hypoamylasemia (d8); AST ↑↑↑ (d8); ALP ↑ (d8); GGT ↑ (d8); CRP ↑↑↑↑ (d8). |
| CYNO1-9 | M | 8 DPI | Decreased appetite (d8); fever (d8). Subject survived to study endpoint (d8). | Lymphocytopenia (d8); thrombocytopenia (d8); monocytopenia (d4, 8); granulocytosis (d4, 8); hypoamylasemia (d8); BUN ↑ (d8); ALT ↑ (d8); AST ↑↑ (d8); ALP ↑ (d8); GGT ↑ (d8). |

Days after BDBV challenge are in parentheses. All reported findings are in comparison to baseline (day of challenge [d0]) values. Decreased appetite is defined as ≤ 65% of food consumed from the previous day. Anorexia is defined as no food consumed from the previous day. Fever is defined as a temperature more than 2.5 °F over baseline, or at least 1.5 °F over baseline and ≥ 103.5 °F. Hypothermia is defined as a temperature ≤3.5°F below baseline. Lymphocytopenia, monocytopenia, erythrocytopenia, thrombocytopenia, and granulocytopenia are defined by a ≥35% drop in numbers of lymphocytes, monocytes, erythrocytes, platelets, or granulocytes (neutrophils, eosinophils, and basophils), respectively. Lymphocytosis, monocytosis, and granulocytosis are defined by a 100% or greater increase in numbers of lymphocytes, monocytes, or granulocytes (neutrophils, eosinophils, and basophils), respectively. Hyperglycemia is defined as a 100% or greater increase in levels of glucose. Hypoglycemia is defined by a ≥25% decrease in levels of glucose. Anemia is defined as a concurrent ≥25% decrease in erythrocyte count, Hct, and Hgb. Hypoalbuminemia is defined by a ≥25% decrease in levels of albumin. Hypoproteinemia is defined by a ≥25% decrease in levels of total protein. Hypoamylasemia is defined by a ≥25% decrease in levels of serum amylase. Hypocalcemia is defined by a ≥25% decrease in levels of serum calcium. Increases in ALT, AST, ALP, CRE, CRP, Hct, and Hgb were graded on the following scale: ↑ = 1-5 fold, ↑↑ = >5-10 fold, ↑↑↑ = >10-20 fold, ↑↑↑↑ = >20-fold, ↓ = ≥50% decrease. (BUN) blood urea nitrogen, (ALT) alanine aminotransferase, (AST) aspartate aminotransferase, (ALP) alkaline phosphatase, (CRE) Creatinine, (CRP) C-reactive protein, (Hct) hematocrit, (Hgb) hemoglobin.

**Supplementary Table 2. Clinical description and outcome of cynomolgus macaques challenged with Bundibugyo virus (BDBV).**

| **Subject No.** | **Sex** | **Clinical illness** | **Clinical pathology** |
| --- | --- | --- | --- |
| CYNO2-1 | M | Fever (d7); anorexia (d8-10); decreased appetite (d11, 12); petechial rash (d8-13); depression (d9-11); hunched posture (d9-11). Subject survived to study endpoint (d28). | Lymphocytopenia (d7); thrombocytopenia (d7, 9); monocytopenia (d1, 5, 7); lymphocytosis (d15); granulocytosis (d5, 7); Hct ↓ (d7); hypoalbuminemia (d15); hypoamylasemia (d7, 9); BUN ↑ (d9); CRE ↑ (d9); ALT ↑ (d7, 12), ↑↑ (d9); AST ↑↑ (d7, 12), ↑↑↑↑ (d9); GGT ↑↑ (d9), ↑ (d12, 15); CRP ↑↑↑ (d7, 9). |
| CYNO2-2 | F | Fever (d7); decreased appetite (d7, 8, 10, 11, 16, 17); anorexia (d9, 12-15); petechial rash (d9, 10, 12, 13); depression (d17); weakness (d17); hunched posture (d17); epistaxis (d17); dyspnea (d17); recumbency (d17). Subject euthanized (d17). | Lymphocytopenia (d7, 9, 12, 15); thrombocytopenia (d7, 9, 12, 15, 17); monocytopenia (d5, 7, 9, 12, 15, 17); granulocytopenia (d17); granulocytosis (d5, 7, 12, 15); hypoglycemia (d7, 12, 15, 17); hypoalbuminemia (d9, 12, 15, 17); hypocalcemia (d17); hypoproteinemia (d17); hypoamylasemia (7, 9, 12, 15); ALT ↑ (d9, 15, 17); AST ↑ (d7), ↑↑↑ (d9), ↑↑ (d12, 15, 17); ALP ↑ (d9, 12, 15, 17); GGT ↑ (d9, 12, 15, 17); CRP ↑↑↑↑ (d7, 9, 15, 17), ↑ (d12). |
| CYNO2-3 | M | Fever (d7); decreased appetite (d7, 12, 13, 22, 24); anorexia (d8-11, 23); depression (d8-14, 23); hunched posture (d8-14, 22-25); hypothermia (d12); diarrhea (d21-28). Subject survived to study endpoint (d28). | Lymphocytopenia (d5, 7, 9); thrombocytopenia (d7, 12); monocytopenia (d5, 9, 28); monocytosis (d15); granulocytosis (d1, 4, 7, 9, 15, 21, 28); hypoalbuminemia (d9, 12, 15, 21, 28); hypoamylasemia (d7, 9, 12); CRE ↑ (d9); ALT ↑ (d9, 12); AST ↑ (d7, 12), ↑↑↑ (d9); GGT ↑ (d9); CRP ↑ (d5), ↑↑↑↑ (d7, 9, 15, 21). |
| CYNO2-4 | F | Decreased appetite (d3-8); fever (d5, 7, 9); anorexia (d9); petechial rash (d7-9); depression (d8, 9); hunched posture (d8, 9); weakness (d9); recumbency (d9); unresponsiveness (d9). Subject euthanized (d9). | Lymphocytopenia (d3, 5, 7); thrombocytopenia (d7, 9); granulocytosis (d1, 5, 7, 9); hypoglycemia (d9); hypoalbuminemia (d9); hypocalcemia (d9); hypoamylasemia (d7, 9); BUN ↑ (d9); CRE ↑ (d9); ALT ↑ (d7), ↑↑↑ (d9); AST ↑↑↑ (d7), ↑↑↑↑ (d9); ALP ↑ (d9); GGT ↑ (d9); CRP ↑ (d5), ↑↑↑↑ (d7). |
| CYNO2-5 | M | Anorexia (d8-10); depression (d8-10); hunched posture (d8-10); petechial rash (d8-10); epistaxis (d10); weakness (d10); ataxia (d10). Subject succumbed to disease (d11). | Lymphocytopenia (d5, 7, 9); thrombocytopenia (d7, 9); monocytopenia (d1, 5, 7, 9); granulocytosis (d3, 5); hypoglycemia (d9); hypoalbuminemia (d9); hypoamylasemia (d7, 9); BUN ↑ (d9); CRE ↑ (d9); ALT ↑↑ (d9); AST ↑ (d1, 7), ↑↑↑ (d9); CRP ↑ (d5), ↑↑↑↑ (d7, 9). |
| CYNO2-6 | M | Inguinal lymphadenomegaly (d5); decreased appetite (d7, 8, 13); anorexia (d9-12, 14-16); petechial rash (d8-16); depression (d9-16); hunched posture (d9-16); diarrhea (d15). Subject succumbed to disease (d16). | Lymphocytopenia (d5, 7, 9); thrombocytopenia (d7, 9, 12); monocytopenia (d1, 12); granulocytosis (d1, 5, 7, 9, 12, 15); anemia (d17); hypoglycemia (d17); hypoalbuminemia (d9, 12, 15, 17); hypoamylasemia (d7, 9, 12); BUN ↑ (d12), ↑↑↑ (d15, 17); CRE ↑ (d12, 15), ↑↑↑ (d17); ALT ↑ (d9, 12, 15, 17); AST ↑ (d7), ↑↑↑↑ (d9, 12, 15, 17); ALP ↑ (d9); CRP ↑↑ (d5), ↑↑↑↑ (d7, 17), ↑ (d9, 12, 15). |
| CYNO2-7 | M | Decreased appetite (d1-5); anorexia (d6-16); depression (d11-16); hunched posture (d11-15); epistaxis (d13, 14, 16); weakness (d16); ataxia (d16); dyspnea (d16); recumbency (d16). Subject euthanized (d16). | Lymphocytopenia (d7, 9, 12); thrombocytopenia (d7, 9, 15); monocytosis (d9); granulocytosis (d3, 5, 7, 9, 12, 15, 16); hypoalbuminemia (d12, 15, 16); hypoamylasemia (d7); BUN ↑ (d12), ↑↑ (d15, 16); CRE ↑ (d1, 3, 5, 7, 9, 12, 15, 16); ALT ↑ (d9, 12, 15, 16); AST ↑ (d1, 7), ↑↑ (d9), ↑↑↑ (d12, 15, 16); ALP ↑ (d9, 12, 15, 16); GGT ↑ (d9, 12, 15, 16); CRP ↑ (d5, 12), ↑↑ (d7, 9, 15, 16). |
| CYNO2-8 | F | Decreased appetite (d6-8, 17-21, 23-28); fever (d7, 9); anorexia (d9-16, 22); depression (d11-17); hunched posture (d11-17). Subject survived to study endpoint (d28). | Lymphocytopenia (d5, 7, 9, 12); thrombocytopenia (d7,9, 12, 15); monocytopenia (d3, 9, 12, 28); monocytosis (d21); granulocytosis (d5, 12, 15, 21, 28); hypoalbuminemia (d9, 12, 15, 21, 28); hypoamylasemia (d7, 9, 12, 15); BUN ↑ (d15); CRE ↑ (d15); ALT ↑ (d7, 15), ↑↑ (d9, 12); AST ↑ (d7, 15), ↑↑↑ (d9, 12); ALP ↑ (7, 9, 21, 28), ↑↑↑ (d12), ↑↑ (d15); GGT ↑ (d9, 15), ↑↑ (d12); CRP ↑ (d5), ↑↑↑ (d7, 9, 12), ↑↑ (d15). |
| CYNO2-9 | F | Decreased appetite (d4, 5, 7, 8); anorexia (d9-15); depression (d9-15); hunched posture (d9-15); rhinorrhea (d12-15); epistaxis (d12-15); petechial rash (d12-15); facial edema (d14, 15); weakness (d15); diarrhea (d15); dyspnea (d15); recumbency (d15); unresponsiveness (d15). Subject euthanized (d15). | Lymphocytopenia (d3, 5, 7, 12); thrombocytopenia (d7, 9, 12, 15); monocytopenia (d5, 7); granulocytopenia (d15); granulocytosis (d3, 5, 7, 9, 12); hypoalbuminemia (d7, 9, 12, 15); hypocalcemia (d15); hypoamylasemia (d7, 9); BUN ↑ (d12), ↑↑ (d15); CRE ↑ (d9, 15); ALT ↑ (d9, 12); AST ↑ (d7, 15), ↑↑↑ (d9, 12); ALP ↑ (d9, 12, 15); GGT ↑ (d9); CRP ↑ (d3), ↑↑ (d7, 9, 12), ↑↑↑↑ (d15). |
| CYNO2-10 | M | Decreased appetite (d6, 7); fever (d7); anorexia (d8-12); depression (d8-12); hunched posture (d8-11); petechial rash (d9-12); ecchymosis (d12); epistaxis (d12); weakness (d12); recumbency (d12); unresponsiveness (d12). Subject euthanized (d12). | Lymphocytopenia (d7, 9); thrombocytopenia (d7, 9, 12); lymphocytosis (d12); monocytosis (d12); granulocytosis (d12); hypoglycemia (d12); hypoalbuminemia (d9, 12); hypoamylasemia (d7, 9); BUN ↑ (d12); CRE ↑ (d12); ALT ↑ (d9), ↑↑ (d12); AST ↑ (d7), ↑↑ (d9), ↑↑↑↑ (d12); ALP ↑ (d9, 12); GGT ↑ (d7), ↑↑ (d9), ↑↑↑ (d12); CRP ↑ (d5, 9), ↑↑↑ (d7, 12). |
| CYNO2-11 | F | Decreased appetite (d1, 3-6, 16-19, 22); anorexia (d7-15); depression (d11-13); hunched posture (d11-13); hypothermia (d21). Subject survived to study endpoint (d28). | Lymphocytopenia (d5, 7, 9, 12, 15); thrombocytopenia (d7, 9, 12, 15); monocytopenia (d1, 3, 5, 7, 9, 12, 15); granulocytopenia (d9, 12, 21, 28); hypoglycemia (d28); hypoalbuminemia (d7, 9, 12, 15, 21); hypoamylasemia (d5, 7, 9); ALT ↑ (d7, 0, 12, 15, 28); AST ↑ (d7, 21), ↑↑ (d9, 12, 15); ALP ↑↑ (d9, 12, 15), ↑ (d21, 28); GGT ↑ (d9, 12, 15); CRP ↑↑ (d5), ↑↑↑ (d7), ↑ (d9, 12). |
| CYNO2-12 | M | Decreased appetite (d6); anorexia (d7-9); depression (d9); weakness (d9); petechial rash (d9); emesis (d9); recumbency (d9). Subject euthanized (d9). | Lymphocytopenia (d5, 7); thrombocytopenia (d5, 7, 9); lymphocytosis (d9); monocytosis (d9); granulocytosis (d3, 5, 9); hypoalbuminemia (d9); hypoamylasemia (d7); BUN ↑ (d9); CRE ↑ (d9); ALT ↑ (d7), ↑↑↑↑ (d9); AST ↑ (d7), ↑↑↑↑ (d9); ALP ↑ (d9); GGT ↑↑ (d9); CRP ↑ (d5), ↑↑ (d7, 9). |

Days after BDBV challenge are in parentheses. All reported findings are in comparison to baseline (day of challenge [d0]) values. Decreased appetite is defined as ≤ 65% of food consumed from the previous day. Anorexia is defined as no food consumed from the previous day. Fever is defined as a temperature more than 2.5 °F over baseline, or at least 1.5 °F over baseline and ≥ 103.5 °F. Hypothermia is defined as a temperature ≤3.5°F below baseline. Lymphocytopenia, monocytopenia, erythrocytopenia, thrombocytopenia, and granulocytopenia are defined by a ≥35% drop in numbers of lymphocytes, monocytes, erythrocytes, platelets, or granulocytes (neutrophils, eosinophils, and basophils), respectively. Lymphocytosis, monocytosis, and granulocytosis are defined by a 100% or greater increase in numbers of lymphocytes, monocytes, or granulocytes (neutrophils, eosinophils, and basophils), respectively. Hyperglycemia is defined as a 100% or greater increase in levels of glucose. Hypoglycemia is defined by a ≥25% decrease in levels of glucose. Anemia is defined as a concurrent ≥25% decrease in erythrocyte count, Hct, and Hgb. Hypoalbuminemia is defined by a ≥25% decrease in levels of albumin. Hypoproteinemia is defined by a ≥25% decrease in levels of total protein. Hypoamylasemia is defined by a ≥25% decrease in levels of serum amylase. Hypocalcemia is defined by a ≥25% decrease in levels of serum calcium. Increases in ALT, AST, ALP, CRE, CRP, Hct, and Hgb were graded on the following scale: ↑ = 1-5 fold, ↑↑ = >5-10 fold, ↑↑↑ = >10-20 fold, ↑↑↑↑ = >20-fold, ↓ = ≥50% decrease. (BUN) blood urea nitrogen, (ALT) alanine aminotransferase, (AST) aspartate aminotransferase, (ALP) alkaline phosphatase, (CRE) Creatinine, (CRP) C-reactive protein, (Hct) hematocrit, (Hgb) hemoglobin.

**Supplementary Table 3. H/E and IHC severity scores of BDBV-challenged cynomolgus macaques from BDBV temporal euthanasia study**

| **Animal ID** | **CYNO1-1** | **CYNO1-2** | **CYNO1-3** |  | **CYNO1-4** | **CYNO1-5** | **CYNO1-6** |  | **CYNO1-7** | **CYNO1-8** | **CYNO1-9** |
| --- | --- | --- | --- | --- | --- | --- | --- | --- | --- | --- | --- |
| **DPI** | 2 | 2 | 2 |  | 4 | 4 | 4 |  | 8 | 8 | 8 |
| **Cardiorespiratory System** |  | | | | | | | | | | |
| Right upper lobe: Representative section | 0 0 | 0 0 | 0 0 |  | 0 0 | 0 0 | 0 0 |  | 1 1 | 1 1 | 1 1 |
| Right middle lobe: Representative section | 0 0 | 0 0 | 0 0 |  | 0 0 | 0 0 | 0 0 |  | 1 1 | 1 1 | 1 1 |
| Right lower lobe: Representative section | 0 0 | 0 0 | 0 0 |  | 0 0 | 0 0 | 0 0 |  | 1 1 | 1 1 | 1 1 |
| Left upper lobe: Representative section | 0 0 | 0 0 | 0 0 |  | 0 0 | 0 0 | 0 0 |  | 1 1 | 1 1 | 1 1 |
| Left middle lobe: Representative section | 0 0 | 0 0 | 0 0 |  | 0 0 | 0 0 | 0 0 |  | 1 1 | 1 1 | 1 1 |
| Left lower lobe: Representative section | 0 0 | 0 0 | 0 0 |  | 0 0 | 0 0 | 0 0 |  | 1 1 | 1 1 | 1 1 |
| Heart | 0 0 | 0 0 | 0 0 |  | 0 0 | 0 0 | 0 0 |  | 0 1 | 0 0 | 0 1 |
| **Lymphoid System** |  | | | | | | | | | | |
| Tonsil | 0 0 | 0 0 | 0 0 |  | 0 0 | 0 0 | 0 0 |  | 2 2 | 0 1 | 0 1 |
| Mandibular lymph node | 0 0 | 0 0 | 0 0 |  | 0 0 | 0 1 | 0 1 |  | 2 2 | 1 1 | 2 2 |
| Axillary lymph node | 0 0 | 0 0 | 0 0 |  | 1 1 | 0 0 | 0 0 |  | 2 2 | 0 0 | 2 2 |
| Inguinal lymph node | 0 0 | 0 0 | 0 0 |  | 1 1 | 1 1 | 0 1 |  | 1 1 | 0 0 | 2 2 |
| Mesenteric lymph node | 0 0 | 0 0 | 0 0 |  | 0 1 | 0 0 | 0 0 |  | 2 2 | 0 0 | 1 1 |
| Spleen | 0 0 | 0 0 | 0 0 |  | 0 1 | 0 1 | 0 1 |  | 3 2 | 2 2 | 3 2 |
| **Urogenital and Endocrine System** |  | | | | | | | | | | |
| Kidney | 0 0 | 0 0 | 0 0 |  | 0 0 | 0 0 | 0 0 |  | 0 1 | 0 0 | 1 1 |
| Urinary Bladder | 0 0 | 0 0 | 0 0 |  | 0 0 | 0 0 | 0 0 |  | 1 2 | 0 0 | 1 1 |
| Gonad | 0 0 | 0 0 | 0 0 |  | 0 0 | 0 0 | 0 0 |  | 1 1 | 0 2 | 1 1 |
| Uterus or Prostate | 0 0 | 0 0 | 0 0 |  | 0 0 | 0 0 | 0 0 |  | 1 1 | 0 2 | 1 2 |
| Adrenal gland | 0 0 | 0 0 | 0 0 |  | 0 0 | 0 0 | 0 0 |  | 0 2 | 0 1 | 1 1 |
| Thyroid gland | 0 0 | 0 0 | 0 0 |  | 0 0 | 0 0 | 0 0 |  | 0 1 | 0 0 | 0 1 |
| **Central Nervous System** |  | | | | | | | | | | |
| Cerebrum-Frontal lobe | 0 0 | 0 0 | 0 0 |  | 0 0 | 0 0 | 0 0 |  | 0 0 | 0 0 | 0 0 |
| Brainstem & Cerebellum | 0 0 | 0 0 | 0 0 |  | 0 0 | 0 0 | 0 0 |  | 0 0 | 0 0 | 0 0 |
| Cerebrum-Temporal & Parietal lobes | 0 0 | 0 0 | 0 0 |  | 0 0 | 0 0 | 0 0 |  | 0 0 | 0 0 | 0 1 |
| Cervical spinal cord | 0 0 | 0 0 | 0 0 |  | 0 0 | 0 0 | 0 0 |  | 0 0 | 0 0 | 0 0 |
| Pituitary gland | 0 0 | 0 0 | 0 0 |  | 0 0 | 0 0 | 0 0 |  | 0 1 | 0 1 | 0 1 |
| **Gastrointestinal System** |  | | | | | | | | | | |
| Liver | 0 0 | 0 0 | 0 0 |  | 0 1 | 1 1 | 0 1 |  | 2 2 | 2 2 | 2 2 |
| Submandibular salivary gland | 0 0 | 0 0 | 0 0 |  | 0 0 | 0 0 | 0 0 |  | 1 1 | 0 0 | 1 1 |
| Pancreas | 0 0 | 0 0 | 0 0 |  | 0 0 | 0 0 | 0 0 |  | 0 1 | 0 0 | 1 1 |
| Stomach | 0 0 | 0 0 | 0 0 |  | 0 0 | 0 0 | 0 1 |  | 1 1 | 0 0 | 1 1 |
| Duodenum | 0 0 | 0 0 | 0 0 |  | 0 0 | 0 0 | 0 1 |  | 2 2 | 1 1 | 1 1 |
| Small intestine-ileum | 0 0 | 0 0 | 0 0 |  | 0 0 | 0 0 | 0 0 |  | 1 2 | 1 1 | 1 1 |
| Cecum | 0 0 | 0 0 | 0 0 |  | 0 0 | 0 0 | 0 0 |  | 2 2 | 1 1 | 1 1 |
| Colon | 0 0 | 0 0 | 0 0 |  | 0 0 | 0 0 | 0 0 |  | 2 2 | 1 1 | 0 1 |
| **Other Systems** |  | | | | | | | | | | |
| Haired Skin/Mammary gland | 0 0 | 0 0 | 0 0 |  | 0 0 | 0 0 | 0 0 |  | 0 1 | 0 0 | 0 1 |
| Nasal mucosa | 0 0 | 0 0 | 0 0 |  | 0 0 | 0 0 | 0 0 |  | 0 0 | 0 0 | 0 0 |
| Conjunctiva/Palpebrae | 0 0 | 0 0 | 0 0 |  | 0 0 | 0 0 | 0 0 |  | 1 2 | 0 0 | 0 1 |
| Eye | 0 0 | 0 0 | 0 0 |  | 0 0 | 0 0 | 0 0 |  | 1 1 | 0 0 | 1 1 |
| **Gender** | M | F | F |  | M | F | F |  | M | F | M |

| ***First score indicates severity score of H&E slide, Second score indicates severity score of IHC slide*** |
| --- |
| 1= Inflammation/lesions in ~10% or less of the examined tissues and IHC labeling |
| 2=Inflammation/lesions in ~25% or less of the examined tissues and IHC labeling |
| 3=Inflammation/lesions in ~50% or less of the examined tissues and IHC labeling |
| 4=Inflammation/lesions in over ~50% of the examined tissues and IHC labeling |
| AKOYA |
| IHC + |

**Supplementary Table 4. H/E and IHC severity scores of BDBV-challenged cynomolgus macaques from BDBV natural history study**

| **Animal ID** | **CYNO2-1** | **CYNO2-2** | **CYNO2-3** | **CYNO2-4** | **CYNO2-5** | **CYNO2-6** | **CYNO2-7** | **CYNO2-8** | **CYNO2-9** | **CYNO2-10** | **CYNO2-11** | **CYNO2-12** |
| --- | --- | --- | --- | --- | --- | --- | --- | --- | --- | --- | --- | --- |
| **DPI** | 28 | 17 | 28 | 9 | 11 | 17 | 16 | 28 | 15 | 12 | 28 | 9 |
| **Cardiorespiratory System** |  |  |  |  |  |  |  |  |  |  |  |  |
| Right upper lobe: Representative section | 0 0 | 0 0 | 0 0 | 2 2 | 1 1 | 0 0 | 0 0 | 0 0 | 0 0 | 2 2 | 0 0 | 2 2 |
| Right middle lobe: Representative section | 0 0 | 0 0 | 0 0 | 2 2 | 1 1 | 0 0 | 0 0 | 0 0 | 0 0 | 2 2 | 0 0 | 2 2 |
| Right lower lobe: Representative section | 0 0 | 0 0 | 0 0 | 2 2 | 1 1 | 0 0 | 0 0 | 0 0 | 1 1 | 2 2 | 0 0 | 2 2 |
| Left upper lobe: Representative section | 0 0 | 0 0 | 0 0 | 2 2 | 1 1 | 0 0 | 0 0 | 0 0 | 1 1 | 2 2 | 0 0 | 2 2 |
| Left middle lobe: Representative section | 0 0 | 0 0 | 0 0 | 2 2 | 2 2 | 0 0 | 0 0 | 0 0 | 1 1 | 2 2 | 0 0 | 2 2 |
| Left lower lobe: Representative section | 0 0 | 0 0 | 0 0 | 2 2 | 2 2 | 0 0 | 0 0 | 0 0 | 1 1 | 2 2 | 0 0 | 2 2 |
| Heart | 0 0 | 0 1 | 0 0 | 0 1 | 0 1 | 0 1 | 0 0 | 0 0 | 0 1 | 0 1 | 0 0 | 0 1 |
| **Lymphoid System** |  |  |  |  |  |  |  |  |  |  |  |  |
| Tonsil | 0 0 | 1 1 | 0 0 | N/A | 3 3 | N/A | 1 1 | 0 0 | 1 1 | 3 2 | 0 0 | 3 2 |
| Mandibular lymph node | 0 0 | 1 1 | 0 0 | 2 2 | 3 2 | 1 1 | 1 1 | 0 0 | 1 1 | 3 1 | 0 0 | 2 2 |
| Axillary lymph node | 0 0 | 1 1 | 0 0 | 2 2 | 3 2 | 1 1 | 1 1 | 0 0 | 1 1 | 3 2 | 0 0 | 3 2 |
| Inguinal lymph node | 0 0 | 1 1 | 0 0 | 3 2 | 3 2 | 1 1 | 2 2 | 0 0 | 1 1 | 3 2 | 0 0 | 3 2 |
| Mesenteric lymph node | 0 0 | 3 2 | 0 0 | 2 2 | 3 2 | 4 1 | 2 2 | 0 0 | 1 1 | 3 2 | 0 0 | 3 2 |
| Spleen | 0 0 | 4 1 | 0 0 | 3 2 | 4 1 | 4 1 | 2 1 | 0 0 | 1 1 | 4 1 | 0 0 | 4 1 |
| **Urogenital and Endocrine System** |  |  |  |  |  |  |  |  |  |  |  |  |
| Kidney | 0 0 | 2 1 | 0 0 | 2 2 | 2 2 | 1 1 | 2 1 | 1 0 | 1 1 | 1 1 | 1 0 | 2 2 |
| Urinary Bladder | 0 0 | 0 0 | 0 0 | 2 2 | 2 2 | 2 2 | 0 0 | 0 0 | 2 2 | 2 2 | 0 0 | 2 3 |
| Gonad | 0 0 | 1 2 | 0 0 | 0 2 | 2 2 | 1 1 | 2 1 | 0 0 | 2 2 | 2 2 | 0 0 | 1 2 |
| Uterus or Prostate | 0 0 | 0 0 | 0 0 | 1 1 | 0 1 | 0 0 | 0 0 | 0 0 | 2 3 | 1 1 | 0 0 | 1 1 |
| Adrenal gland | 0 0 | 2 2 | 0 0 | 1 2 | 1 2 | 2 2 | 1 1 | 0 0 | 1 1 | 2 2 | 0 0 | 1 2 |
| Thyroid gland | 0 0 | 1 1 | 0 0 | 0 1 | 1 2 | 0 1 | 1 1 | 0 0 | 2 2 | 1 1 | 0 0 | 1 2 |
| **Central Nervous System** |  |  |  |  |  |  |  |  |  |  |  |  |
| Cerebrum-Frontal lobe | 0 0 | 0 0 | 0 0 | 0 0 | 0 0 | 0 0 | 0 0 | 0 0 | 0 0 | 0 0 | 0 0 | 0 0 |
| Brainstem | 0 0 | 2 2 | 0 0 | 1 1 | 1 1 | 2 2 | 1 1 | 0 2 | 1 1 | 1 1 | 0 0 | 2 2 |
| Cerebellum | 0 0 | 0 0 | 0 0 | 0 0 | 0 0 | 0 0 | 0 0 | 0 0 | 0 0 | 0 0 | 0 0 | 0 0 |
| Cerebrum-Temporal & Parietal lobes | 0 0 | 0 0 | 0 0 | 1 1 | 1 1 | 2 2 | 1 1 | 0 0 | 1 1 | 1 1 | 0 0 | 2 2 |
| Cervical spinal cord | 0 0 | 0 0 | 0 0 | 0 0 | 0 0 | 0 0 | 0 0 | 0 0 | 0 0 | 0 0 | 0 0 | 0 0 |
| Pituitary gland | 0 0 | 1 1 | 0 0 | 1 1 | 1 1 | 2 2 | 2 2 | 0 0 | 2 2 | 1 1 | 0 0 | 2 2 |
| **Gastrointestinal System** |  |  |  |  |  |  |  |  |  |  |  |  |
| Liver | 1 0 | 3 1 | 0 0 | 3 3 | 3 1 | 2 0 | 1 1 | 1 0 | 2 0 | 3 2 | 0 0 | 3 3 |
| Submandibular salivary gland | 0 0 | 0 0 | 0 0 | 1 1 | 0 0 | 0 0 | 0 0 | 0 0 | 0 0 | 0 0 | 0 0 | 1 1 |
| Pancreas | 0 0 | 3 3 | 0 0 | 1 2 | 1 2 | 3 3 | 3 3 | 0 0 | 3 3 | 1 2 | 0 0 | 1 2 |
| Tongue | 0 0 | 0 0 | N/A | 1 3 | 1 3 | 0 0 | 0 0 | 0 0 | 0 0 | 1 3 | 0 0 | 1 3 |
| Stomach | 0 0 | 2 2 | 0 0 | 2 2 | 2 2 | 2 2 | 2 2 | 0 0 | 2 2 | 2 2 | 0 0 | 2 2 |
| Duodenum | 0 0 | 2 2 | 0 0 | 2 2 | 2 2 | 2 2 | 2 2 | 0 0 | 0 0 | 2 2 | 0 0 | 2 2 |
| Small intestine-ileum | 0 0 | 2 2 | 0 0 | 2 2 | 2 2 | 2 2 | 2 2 | 0 0 | 0 0 | 2 2 | 0 0 | 2 2 |
| Cecum | 0 0 | 2 2 | 0 0 | 2 2 | 2 2 | 2 2 | 2 2 | 0 0 | 0 0 | 2 2 | 0 0 | 2 2 |
| Colon | 0 0 | 2 2 | 0 0 | 2 2 | 2 2 | 2 2 | 2 2 | 0 0 | 1 1 | 2 2 | 0 0 | 2 2 |
| **Other Systems** |  |  |  |  |  |  |  |  |  |  |  |  |
| Haired Skin/Mammary gland | 0 0 | 2 2 | 0 0 | 1 2 | 0 0 | N/A | 3 3 | N/A | 1 1 | 1 1 | 0 0 | 1 2 |
| Nasal mucosa | 0 0 | 1 2 | 0 0 | 1 1 | 1 1 | 1 1 | 3 3 | 0 0 | 0 0 | 1 1 | 0 0 | 1 2 |
| Conjunctiva/Palpebrae | 0 0 | 1 1 | 0 0 | 1 1 | 1 1 | 1 1 | 1 3 | 0 0 | 1 1 | 1 1 | 0 0 | 2 3 |
| Eye | 0 0 | 0 0 | 0 0 | 1 1 | 1 1 | 1 1 | 2 3 | 0 0 | 2 3 | 1 2 | 0 0 | 1 2 |
| **Gender** | M | F | M | F | M | M | M | F | F | M | F | M |

| ***First score indicates severity score of H&E slide, Second score indicates severity score of IHC slide*** |
| --- |
| 1= Inflammation/lesions in ~10% or less of the examined tissues and IHC labeling |
| 2=Inflammation/lesions in ~25% or less of the examined tissues and IHC labeling |
| 3=Inflammation/lesions in ~50% or less of the examined tissues and IHC labeling |
| 4=Inflammation/lesions in over ~50% of the examined tissues and IHC labeling |
| AKOYA |
| IHC + |

**Supplementary Table 5. Antibodies used for multiplex immunofluorescence (mIF) using the Akoya Biosciences PhenoCycler-Fusion 2.0 platform**

| **Protein specificty** | **Host (clone or lot#)** | **Vendor** | **Dilution** | **Exposure** | **Barcode (Fluorophore)** |
| --- | --- | --- | --- | --- | --- |
| Bundibugyo virus glycoprotein (BDBV) | Rabbit (1306002) | Integrated Biotherapeutics (IBT) | 1:200 | 200ms | BX045 (AF646) |
| CD1d | Mouse (NOR3.2 (NOR3.2/13.17)) | Abcam | 1:200 | 200ms | BX027 (AF646) |
| Macrosialin (CD68) | Rabbit (D4B9C) | Cell Signaling Technology (CST) | 1:200 | 200ms | BX106 (ATTO550) |
| Fibrin β-chain IgG (Fibrin) | Mouse | BioMedica Diagnostics | 1:200 | 200ms | BX003 (AF646) |
| Dendritic cell–specific ICAM-3–grabbing nonintegrin/DC-SIGN (CD209) | Mouse (DCN46) | BD Pharmingen | 1:100 | 500ms | BX031 (AF646) |
| Myeloperoxidase (MPO) | Rabbit (E1E7I) | CST | 1:200 | 200ms | BX015 (AF647) |
| Claudin-5 | Rabbit (EPR7583) | Abcam | 1:200 | 200ms | BX005 (ATTO550) |
| human leukocyte antigen - DR isotype (HLA-DR) | AKYP0063 | Akoya Biosciences/Quanterix | 1:200 | 200ms | BX033 (AF750) |
| B-cell lymphoma 2 (BCL-2) | AKYP0120 | Akoya Biosciences/Quanterix | 1:200 | 200ms | BX085 (AF647) |
| MRP-14 or calgranulin-B (S100A9) | Rabbit (D5060) | CST | 1:200 | 200ms | BX014 (ATTO550) |
| ionized calcium-binding adaptor molecule-1 (IBA-1) | Rabbit (EPR16588) | Abcam | 1:200 | 200ms | BX007 (AF750) |
| glial fibrillary acidic protein (GFAP) | Mouse (SMI24) | BioLegend | 1:200 | 200ms | BX007 (AF750) |
| vimentin | AKYP0082 | Akoya Biosciences/Quanterix | 1:200 | 200ms | BX022 (AF750) |
| aquaporin-4 (AQP4) | Rabbit (D1F8E) | CST | 1:200 | 200ms | BX025 (ATTO550) |
| neuronal nuclei (NeuN) | Mouse (EPR12763) | Abcam | 1:200 | 200ms | BX016 (AF750) |
| Microtubule-Associated Protein 2 (MAP-2) | Rabbit (EPR19691) | Abcam | 1:200 | 200ms | BX010 (AF647) |
| Oligodendrocyte transcription factor 2 (Olig2) | Rabbit (EPR2673) | Abcam | 1:200 | 200ms | BX024 (AF647) |
